# Regulation of terminal hypertrophic chondrocyte differentiation in *Prmt5* mutant mice modeling infantile idiopathic scoliosis

**DOI:** 10.1101/683136

**Authors:** Zhaoyang Liu, Janani Ramachandran, Steven A Vokes, Ryan S Gray

**Affiliations:** Department of Pediatrics, Dell Pediatric Research Institute, University of Texas at Austin Dell Medical School, 1400 Barbara Jordan Blvd., Austin, TX 78723 USA; Department of Molecular Biosciences, University of Texas at Austin, 100 E 24^th^ Street Stop A5000, Austin, TX 78712 USA; Department of Nutritional Sciences, University of Texas at Austin, 103 W. 24^th^ Street A2703, Austin, TX 78712 USA

**Keywords:** Hypertrophic chondrocyte differentiation, skeletogenesis, Scoliosis

## Abstract

Idiopathic scoliosis (IS) is the most common type of musculoskeletal defect effecting children and is classified by age of onset, location, and degree of spine curvature. Although rare, the onset of IS during infancy is the more severe and rapidly progressive form of the disease, leading to increased mortality due to significant respiratory compromise. The pathophysiology of IS, in particular for infantile IS, remain elusive. Here, we show that PRMT5 is critical for the regulation of terminal hypertrophic chondrocyte differentiation in the spine and models infantile IS in mouse. Conditional ablation of PRMT5 in osteochondral progenitors led to impaired terminal hypertrophic chondrocyte differentiation and asymmetric defects of endochondral bone formation in the perinatal spine. Analysis of several markers of endochondral ossification revealed increased COLX and *Ihh* expression and a dramatic reduction of *Mmp13* and RUNX2 expression in the intervertebral disc and vertebral growth plate. Furthermore, we demonstrate that PRMT5 function in committed chondrogenic lineages is required for regulation of COLX expression in the adult spine. Together, our results establish PRMT5 as a critical regulator of hypertrophic chondrocyte differentiation and endochondral bone formation in spine development and maintenance.

## INTRODUCTION

Idiopathic scoliosis (IS) is the most common pediatric spinal deformity characterized by a lateral curvature of the spine >10°, affecting ∼3% of children worldwide (Wise et al., 2008). Clinical sub-classifications of IS are based on age of presentation at infantile (birth to 3 years), juvenile (age 3 to 11 years), and adolescent (11 years and older) ages (Giampietro et al., 2003). Infantile IS develops rapidly and can lead to significant respiratory compromise, increased morbidity, and mortality (Cahill and Samdani, 2012; Davies and Reid, 1970; Pehrsson et al., 1992; Roye, 2018). Therefore, there is a strong need for early diagnosis and rational therapies that might halt or ameliorate the pathogenesis of infantile IS.

The etiology of IS remains poorly understood, however a genetic causality and candidate loci associated with the disease are beginning to be established. For instance, sibling risk studies report that 11.5-19% of siblings share risk for spine curvatures >10° (Riseborough and Wynne-Davies, 1973; Rogala et al., 1978). Large-scale genome wide association studies also implicated several candidate loci associated with IS, including associations with *GPR126* (*ADGRG6*), *LBX1*, *CHL1* and *SOX9* genes (Ikegawa, 2016; Kou et al., 2013; Sharma et al., 2011; Takahashi et al., 2011). Some of these candidate genes associated with IS and their effectors, including *Adgrg6/Gpr126* (Karner et al., 2015), *Sox9* (Henry et al., 2012), *Shp2* (Kim et al., 2013), *Gdf5/6* (Settle et al., 2003), and *Fgf3* (Gao et al., 2015), are known to develop spine defects and scoliosis when mutated in mouse. Interestingly, a majority of these genes are involved with the development and homeostasis of connective tissues and cartilages, indicating a potential link between cartilaginous tissues and the pathogenesis of IS (Liu and Gray, 2018).

The intervertebral disc (IVD) is a cartilaginous tissue that connects each of the vertebral bodies in the spine, functioning to both disperse mechanical loading and convey flexibility to the spine. The IVD is composed of the nucleus pulposus, the inner most gel-like center, which is surrounded by numerous fibrocartilage lamellar layers collectively known as the annulus fibrosus. The IVD is connected to the growth plate and bony vertebrae by the cartilaginous endplate. The vertebrae form via endochondral ossification of the cartilaginous anlage flanking the notochord during embryonic development and elongate through proliferation and differentiation of the growth plate during postnatal development (Smith et al., 2011).

Endochondral ossification is the process by which the majority of the mammalian skeleton is formed. It is a complex process in the vertebral bodies, which is comparable to homologous mechanisms of that in long bone (Karsenty et al., 2009; Long and Ornitz, 2013). This process is characterized by the succession of proliferative, prehypertrophic, hypertrophic, and terminal hypertrophic chondrocytes (Long and Ornitz, 2013). The ossification of the vertebra begins within the center of the cartilaginous vertebral body template. Cells within this center of ossification exit the cell cycle and initiate hypertrophy associated with the secretion of type X collagen (COLX; encoded by *Col10a1*). The terminal hypertrophic chondrocytes express the matrix metalloproteinase 13 (*Mmp13*), which is critical for terminal differentiation of hypertrophic chondrocytes and endochondral bone formation (Inada et al., 2004). Mechanistically, MMP-13 aids in the degradation of the COLX rich extracellular matrix surrounding the hypertrophic chondrocytes, while invading blood vessels attracted by vascular endothelial growth factors (VEGFs) to promote the infiltration of bone forming osteoblasts (Karsenty et al., 2009; Long and Ornitz, 2013). While many hypertrophic chondrocytes undergo apoptosis clearing a way for the elaboration of the bony matrix, recent studies demonstrate that a subset of these hypertrophic chondrocytes can transdifferentiate into osteoblasts, directly contributing to bone formation (Jing et al., 2015; Park et al., 2015; Zhou et al., 2014).

Chondrocyte maturation and growth plate development are tightly regulated by a series of growth factors and transcriptional regulators. For example, the transcriptional regulator SOX9 (SRY-related high mobility group-box 9) is required for chondrocyte proliferation and to prevent precocious hypertrophic differentiation (Akiyama et al., 2002; Karsenty et al., 2009). Whereas the transcription factor RUNX2 (Runt related transcription factor 2), which has been shown to directly regulate *Col10a1*, *Mmp13* and *Vegfa* expression (Hess et al., 2001; Li et al., 2011; Takahashi et al., 2017; Zelzer et al., 2001), is an important driver of hypertrophic chondrocyte differentiation and endochondral ossification (Komori, 2010a; Komori, 2010b; Komori, 2018). Indian hedgehog (*Ihh*) signaling plays a critical role in chondrocyte proliferation and hypertrophic differentiation via PTHrP dependent and independent processes (Amano et al., 2014; Long and Ornitz, 2013; Mak et al., 2008).

The protein arginine methyltransferase 5 (PRMT5) is required for the maintenance of chondrocyte progenitors in embryonic limb buds (Norrie et al., 2016), where its loss results in precocious *Sox9* expression followed by widespread apoptosis (Norrie et al., 2016). Mechanistically, PRMT5 mediates symmetric demethylation of arginine residues on histones H3R8, H3R2, and H4R3, which regulates a diverse set of target proteins (Bedford and Clarke, 2009; Pal et al., 2004; Stopa et al., 2015; Zhao et al., 2009). PRMT5 dependent arginine methylation can also regulate the assembly of the small nuclear ribonucleoprotein (snRNP) complex (Chari et al., 2008; Meister and Fischer, 2002; Stopa et al., 2015), effecting splicing of mRNAs and gene expression. Recent work also demonstrated that PRMT5 dependent posttranslational di-methlyation of SOX9 protein can increase its half-life (Sun et al., 2019). Given the range of possible molecular functions for PRMT5, it is not surprising that this enzyme plays a role in a number of tissue-specific differentiation pathways, including keratinocytes, muscle, nerve cells and lung differentiation (Calabretta et al., 2018; Chittka et al., 2012; Dacwag et al., 2009; Kanade and Eckert, 2012; Li et al., 2018). In this study, we used conditional mouse genetics coupled with histological and gene expression analyses to establish a novel role for PRMT5 during chondrocyte differentiation for perinatal spine development, modeling infantile IS. We additionally demonstrate a continuous role for PRMT5 in growth plate of the IVD in adult mice. Overall, our work establishes PRMT5 as a critical factor for the terminal differentiation of hypertrophic chondrocytes and endochondral ossification of the spine.

## MATERIALS AND METHODS

### Mice

Animal studies were approved by the Institutional Animal Care and Use Committee at the University of Texas at Austin (AUP-2018-00276 and AUP-2016-00255). All mouse strains, including *Prmt5^f/f^* (Prmt5^tm2c(EUCOMM)Wtsi)^ (Norrie et al., 2016), *Col2Cre* (Long et al., 2001), *ATC* (Dy et al., 2012), *OcCre* (Zhang et al., 2002), and *Rosa-LacZ* (Soriano, 1999), were described previously. Doxycycline (Dox) was administered to *ATC; Prmt5^f/f^* and littermate controls or *ATC; Rosa-LacZ^f/+^* mice with intraperitoneal (IP) injections by two strategies: (i) starting at 2-weeks of age, once/week (10mg Dox/kg body weight) for 4 continuous weeks, or (ii) starting at 4-weeks of age, once/week (10mg Dox/kg body weight) for 4 continuous weeks. Mice were harvested at P1, P10, 6-weeks, 2.5-months and 4-months of age for tissue analysis.

### Analyses of mice

Skeletal preparations were performed as previously described (Allen et al., 2011). Radiographs of mouse skeleton were generated using a Kubtec DIGIMUS X-ray system (Kubtec T0081B) with auto exposure under 25 kV. Cobb angle was measured on high resolution X-ray images with ImageJ as previously described (Cobb, 1948). Histological analysis was performed on thoracic spines fixed in 10% neutral-buffered formalin for 3 days at room temperature followed by 1-week decalcification in Formic Acid Bone Decalcifier (Immunocal, StatLab). After decalcification, bones were embedded in paraffin and sectioned at 5μm thickness. Alcian Blue Hematoxylin/Orange G (ABH/OG) and Safranin O/Fast Green (SO/FG) staining were performed following standard protocols. Quantification of IVD clefts and cell layers were performed on ABH/OG stained sections of 3 control and 3 mutant mice. 3-6 IVDs were analyzed per mouse. Immunohistochemical analyses were performed on paraffin sections after antigen retrieval using 10mM Tris and 1mM EDTA (with 0.05% Triton-X-100, pH 9.0) (PRMT5 and RUNX2), 4mg/ml pepsin (COLII and COLX), 100μg/ml hyaluronidase (PRG4), or 10μg/ml Proteinase K (SOX9), and colorimetric development methodologies with the following primary antibodies: anti-PRMT5 (Abcam, ab109451, 1:100), anti-Collagen Type II (Thermo Scientific, MS235B, 1:100), anti-Collagen Type X (Quartett, 1-CO097-05, 1:200), anti-SOX9 (Santa Cruz Biotechnology, sc-20095, 1:50), anti-Lubricin (PRG4) (Abcam, ab28484, 1:400), and anti-RUNX2 (Medical & Biological Laboratories, D130-3, 1:100). The Terminal deoxynucleotidyl transferase dUTP Nick-End Labeling (TUNEL) cell death assay was performed on paraffin sections with the In Situ Cell Death Detection Kit, Fluorescein (Roche) according to the manufacturer’s instructions. Quantification of TUNEL positive cells were performed on sections of 3 control and 3 mutant mice. 2-4 IVDs were analyzed per mouse. The beta-galactosidase staining was performed on frozen sections as previously described (Liu et al., 2015). Briefly, spines were harvested and fixed in 4% paraformaldehyde for 2 hours at 4 °C and decalcified with 14% EDTA at 4 °C for 1 week. Tissues were washed in sucrose gradient, embedded with Tissue-Tek OCT medium, snap-frozen in liquid nitrogen, and sectioned at 10μm with a Thermo Scientific HM 550 cryostat. *In situ* hybridization using a Digoxygenin-labeled antisense riboprobes for *Mmp13*, *Ihh*, and *Bmp4* were performed on 5μm paraffin sections as described previously with modifications (Karner et al., 2015), and detected with a tyramine-amplified fluorescent antibody (Perkin Elmer, NEL753001KT).

### RNA isolation and Real-time RT-PCR

Intervertebral discs from the thoracic spine of the 2.5-month *ATC; Prmt5^f/f^* and control mice (Dox induced from 4-weeks) were isolated in cold PBS, pooled together, snap frozen and pulverized in liquid nitrogen. Three control and three mutant mice were used in each group. Total RNA was isolated using the TRlzol Reagent (Invitrogen, 15596026), and cleaned up with the Direct-zol RNA miniprep kit (Zymo Research, Z2070). Reverse transcription was performed using 500ng total RNA with the iScript cDNA synthesis kit (BioRad). Real-time RT-PCR analyses were performed as previously described (Liu et al., 2015). Gene expression was normalized to b-actin mRNA and relative expression was calculated using the 2^-(ΔΔCt)^ method. Primers sequences are listed in Table S1.

## RESULTS

### Conditional loss of *Prmt5* in the spine models infantile IS in mouse

In order to test the role of PRMT5 in the spine we crossed mice harboring a *Prmt5^f/f^* conditional allele (Norrie et al., 2016) to *Collagen2al*-Cre (*Col2Cre*) mice (Long et al., 2001). This *Col2Cre* allele demonstrates robust Cre activity in osteochondral progenitors (OCPs) and effectively labeling most structural elements of the spine including the entire IVD, the periosteum, and trabecular bone (Supplemental Fig. *1*). *Col2Cre;Prmt5^f/f^* mutant pups were produced at mendelian ratios when assessed at E18.5 (22.2%, *n=*18). However, all mutant animals died of unknown causes prior to weaning (11.9% survival assessed at Postnatal (P) day 1, *n=*42; and 5.4% survival assessed at P10, *n=*62) (Fig. *1A*). We did not recover any *Col2Cre;Prmt5^f/f^* mutants after P14 (0%, n=23). Two of the mutant mice that survived at P10 (*n*=4) were much smaller than the control mice (Fig. *1F*), while the other two were comparable to the littermate controls, suggesting variable penetrance. Interestingly at E18.5, analysis by whole-mount skeletal prep revealed no obvious defects in the size, patterning, and maturation of the skeleton in the *Col2Cre;Prmt5^f/f^* mutants (Fig. *1C*; *n=*4). However, at P10 we observed obvious IS-like thoracic scoliosis (red arrowhead; Fig. *1F*) with an average cobb angle of 30±4° in *Col2Cre;Prmt5^f/f^* mutants (Fig. *1D*; *n=4/4*). Lateral X-rays had loss of signal attenuation in the distal ribs indicative of reduced ossification in mutant mice (Fig. *1F”*) relative to littermate controls (yellow arrowheads; Fig. *1E”*). Immunohistochemistry (IHC) against PRMT5 revealed robust protein expression throughout the IVD and growth plate in wild-type mice at P1 and P10, which was consistently reduced in *Col2Cre;Prmt5^f/f^* mutant mice (Fig. *1G-J’*). We did not observe PRMT5 expression in the trabecular bone or cortical bone (Supplemental Fig. *2*), suggesting that PRMT5 has a limited role in committed osteoblast lineages.

**Figure 1.**
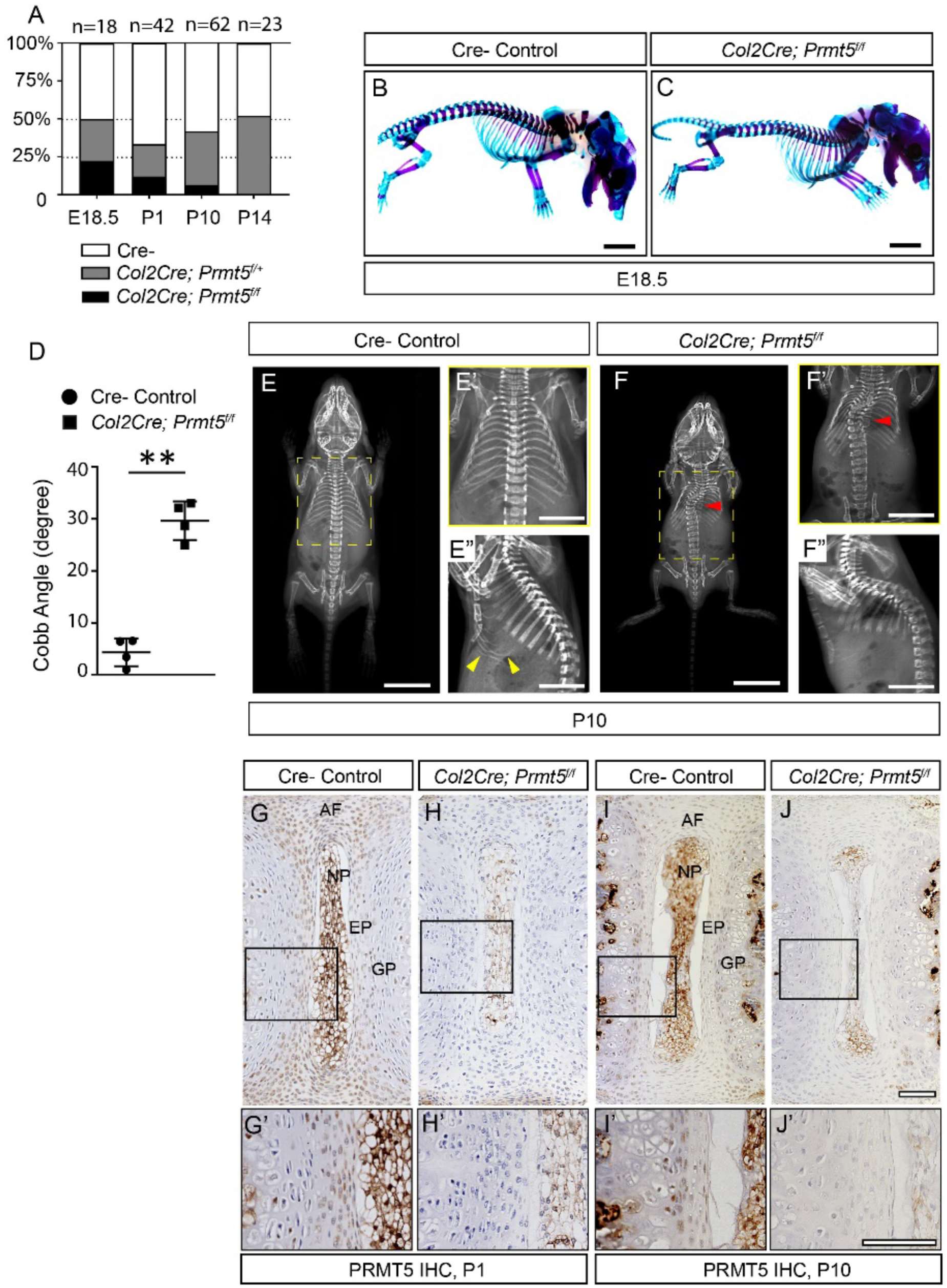
Loss of *Prmt5* in osteochondral progenitor cells induces early onset scoliosis in mice. **(A)** *Col2Cre;Prmt5^f/f^* mutant mice were produced in Mendelian ratios at E18.5 (22.2%, *n*=18), but displayed progressive mortality prior to weaning (11.9%, *n*=42 at P1,5.4%, *n*=62 at P10, and 0%, n=23 at P14). **(B, C)** Skeletal preparations showed comparable size and normal patterning of the spine in *Col2Cre;Prmt5^f/f^* mutant mice at E18.5 compared with the littermate Cre- controls. (*n*=4 for each group.) **(D-F)** X-ray imaging analysis and quantification at P10 demonstrating mutant mice with early-onset thoracic scoliosis (red arrowhead; E and E’) and increased cobb angle (D) (*n*=4 for each group; Each dot represents one mouse analyzed, plotting with mean ± SD; “**” indicates p<0.01, two-tailed student *t*-test). Sagittal X-ray image revealed deficient ossification of the distal ribs in the mutant (F”) compared with a littermate control (yellow arrowheads; E”). **(G-J)** Immunohistochemistry of PRMT5 on both control (G, I) and mutant (H, J) IVD at P1 and P10, respectively. Images with higher magnification were indicated with black boxes and shown in G’, H’, I’ and J’. Strong PRMT5 signal was detected in the entire IVD of the control mice (G, I), but was greatly diminished in the IVDs of *Col2Cre;Prmt5^f/f^* mutant mice (H, J). Scale bars: 2mm in (B, C); 10mm in (E, F); 50mm in (E’-E’’, F’-F’’); and 100μm in (G-J’). *AF: annulus fibrosus, EP: endplate, GP: growth plate, NP: nucleus pulposus*.

### Loss of *Prmt5* in osteochondral progenitors results in asymmetric defects of endochondral ossification in the spine

Using histological approaches, we observed impaired endochondral ossification of the vertebral body in *Col2Cre;Prmt5^f/f^* mutant mice at P1 (Fig. *2B*). In addition, mutant IVDs show an increase of midline clefts in the endplate and growth plate (Fig. *2B’, B”*, quantified in *E*), possibly resulting from the failure of midline fusion during the development of the vertebral bodies and the IVDs (Smith et al., 2011). At P10, there is impaired endochondral ossification and reduced trabecular and cortical bone formation in the vertebral body in a highly heterogeneous manner (Fig. *2D, D”*); in some instances, with stark asymmetries of endochondral ossification even in adjacent vertebrae (yellow asterisks; Fig. *2D*). These regions of persistent cartilage tissues in the vertebrae displayed a general disorganization of the chondrocytes, some of which form into columns or clusters (red, yellow dashed outlines; Fig. *2 D”*). These asymmetrical defects in endochondral ossification are likely to translate to anisotropic mechanical properties of the spine, which in turn may drive the onset of scoliosis observed in *Col2Cre;Prmt5^f/f^* mutant mice.

**Figure 2.**
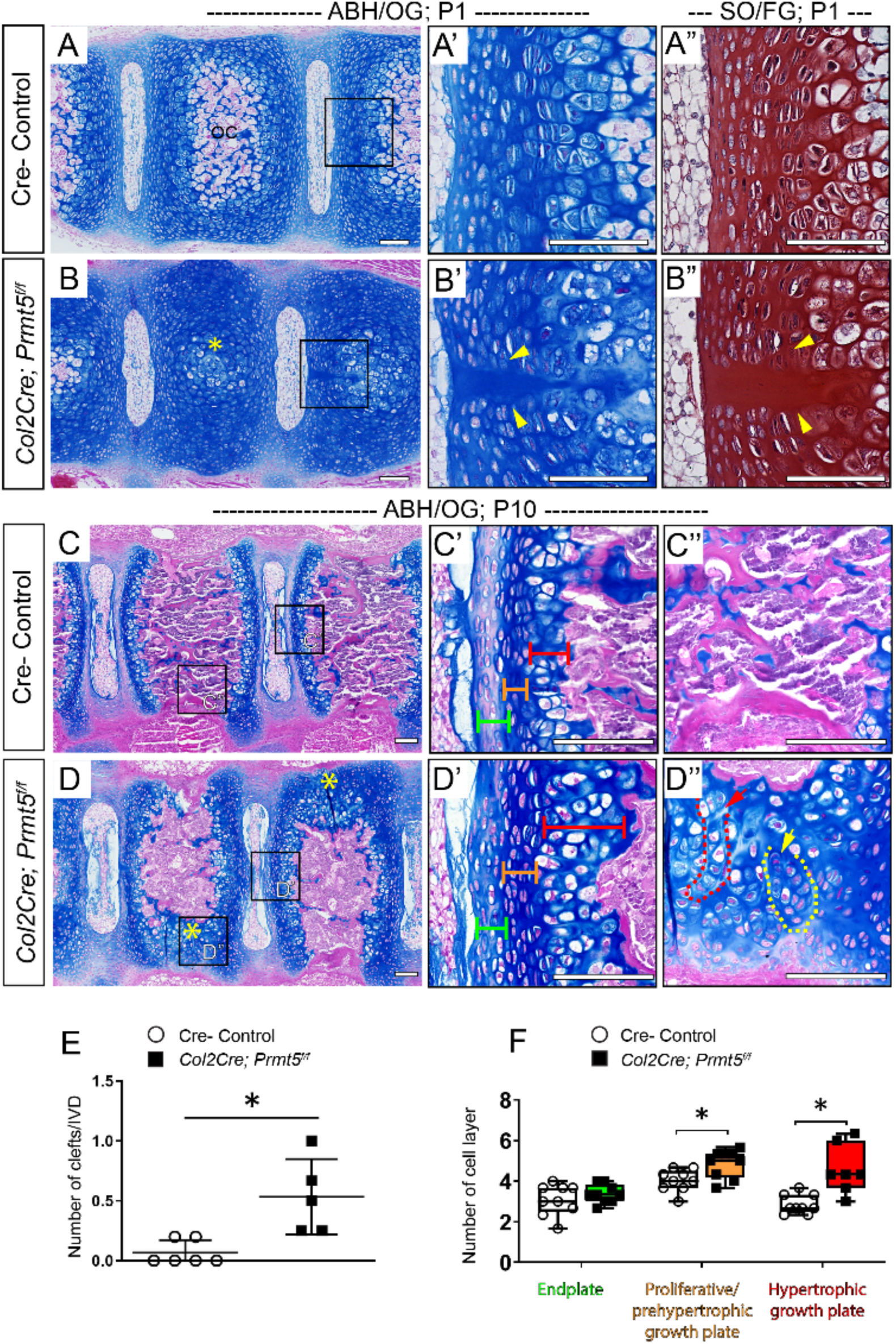
Loss of *Prmt5* in osteochondral progenitor lineages of the spine results in defective ossification of the vertebrae body. **(A-D)** Alcian Blue Hematoxylin/ Orange G (ABH/OG) staining of mouse spine of Cre - control (A and C) or *Col2Cre;Prmt5^f/f^* mutant (B and D) mice at P1 (A and B) and P10 (C and D). Images with higher magnification are indicated with black boxes and shown in (A’-D”). (A” and B”) P1 adjacent sections stained with Safranin O/Fast Green (SO/FG). (B) Impaired ossification in *Col2Cre;Prmt5^f/f^* mutant vertebra (yellow asterisk). (B’, B”) Representative images of midline clefts consistently observed in mutant endplate, and not in Cre -control mice (A’, A”) (yellow arrowheads; *n*=6 for controls and *n*=5 for mutants.) (D) At P10, asymmetric defects of trabecular bone formation were consistently observed in the vertebra of mutant mice. Areas normally ossified in Cre -control mice (C) displayed persistent cartilage in *Col2Cre;Prmt5^f/f^* mutant mice (yellow asterisks; D). The organization of chondrocytes in these regions of persist cartilage were found in columns (D”, red arrow/outline) and clusters (D”, yellow arrow/outline). (*n*=3 for each group.) Endplate, proliferative/prehypertrophic growth plate, and hypertrophic growth plate are labeled with green, orange, and red brackets, respectively. **(E)** Quantification of the increased midline clefts of endplate observed in mutant IVD at P1 as shown in B-B”. Each dot represents one mouse analyzed, plotting with mean ± SD. (*n*=6 mice for controls and *n*=5 mice for mutants.) **(F)** Quantification of the number of cell layers of the endplate and growth plate in control and mutant IVD at P10 as shown in C’ and D’. Each dot in the box and whisker plot represents the counts from a single IVD analyzed. Each IVD was analyzed three times and averaged, and at least two IVDs were analyzed per mouse. (*n*=3 mice for each group.) For all statistical analysis, “*” indicates p<0.05, two-tailed student *t*-test. Scale bars: 100μm in (A-D”).

We also observed morphological alterations in the mutant IVD at P10. In controls, the IVD endplate contains several layers of flat cells that lightly stain with Alcian blue (green bracket; Fig. *2C’*). The proliferative/prehypertrophic zone of the control growth plate is made up of several layers of flat chondrocytes that organize into columns (orange bracket; Fig. *2C’*), while the hypertrophic zone of the growth plate is composed of 2-4 layers of enlarged hypertrophic chondrocytes (red bracket, Fig *2C’*). In *Col2Cre;Prmt5^f/f^* mutant mice we observed alterations of these regions of the IVD. For instance, the cells of the endplate appeared larger, and the surrounding matrix was more heavily stained by Alcian blue (green bracket; Fig. *2D’*), suggesting inappropriate differentiation of the endplate tissue. Quantitative analysis in these IVD tissues demonstrated increased number of cell layers in the hypertrophic zone of the growth plate, but not in the endplate of the *Col2Cre;Prmt5^f/f^* mutant mice (Fig. *2F*). We also observed a mild increased of cell layers in proliferative/prehypertrophic zone (Fig. *2F*). Collectively, these data suggest a model in which PRMT5 is critical for development of vertebra formation due to defects in hypertrophic chondrocyte differentiation.

### Deletion of *Prmt5* in osteochondral progenitors results in altered extracellular matrix components and accumulation of type X collagen in the cartilaginous tissues of the spine

We next assayed several established regulators of hypertrophic chondrocyte differentiation(Long and Ornitz, 2013) in order to assess potential mechanisms for PRMT5-dependent regulation of IVD development. IHC analysis of the hypertrophic chondrocyte marker type X collagen (COLX) revealed ectopic, expanded expression throughout the entire IVD in *Col2Cre;Prmt5^f/f^* mutant mice, and highlighted the expansion of the hypertrophic zone of the growth plate (Fig. *3B, B’*), compared to controls (Fig. *3C, C’*). On the other hand, the expression of proteoglycan 4 (PRG4/ Lubricin), a common marker of healthy IVD (Jay, 1992; Jay and Waller, 2014; Teeple et al., 2015), was remarkably absent in the IVDs of *Col2Cre;Prmt5^f/f^* mutant mice at P10 (Fig. *3D, D’*). The expression of SOX9, a key transcriptional regulator of chondrogenesis, was reduced in the cartilaginous endplate of mutant mice (Supplemental Fig. *3C-D’*), but was not obviously affected in the proliferative growth plate. Hypertrophic chondrocytes of the vertebral growth plate downregulate *Sox9* expression during the process of terminal hypertrophic chondrocyte differentiation (Supplemental Fig. *3C’*). However, *Col2Cre;Prmt5^f/f^* mutant mice displayed ectopic SOX9 expression in these phenotypically hypertrophic cells of the growth plate (Supplemental Fig. *3D’*). Despite these transcriptional alterations of Sox9, we found that type II Collagen (COLII) expression was not obviously changed in *Col2Cre;Prmt5^f/f^* mutant IVDs at P10 (Supplemental Fig. *3A-B’*). Next, we performed the same IHC analysis in the vertebral body of the *Col2Cre;Prmt5^f/f^* mutant mice at P10 as shown in (Fig. *2D, D”*). Within these areas of persistent cartilage in the vertebral bodies, we observed expression of SOX9 and COLX but not PRG4 expression (Supplemental Fig. *4*), suggesting these persistent areas of cartilage in the vertebral bone were the result of similar alterations of hypertrophic chondrocyte differentiation as was observed in the vertebral growth plate. Together our results suggest that PRMT5 regulates the normal development of hypertrophic chondrocytes of the growth plate and vertebral bodies.

**Figure 3.**
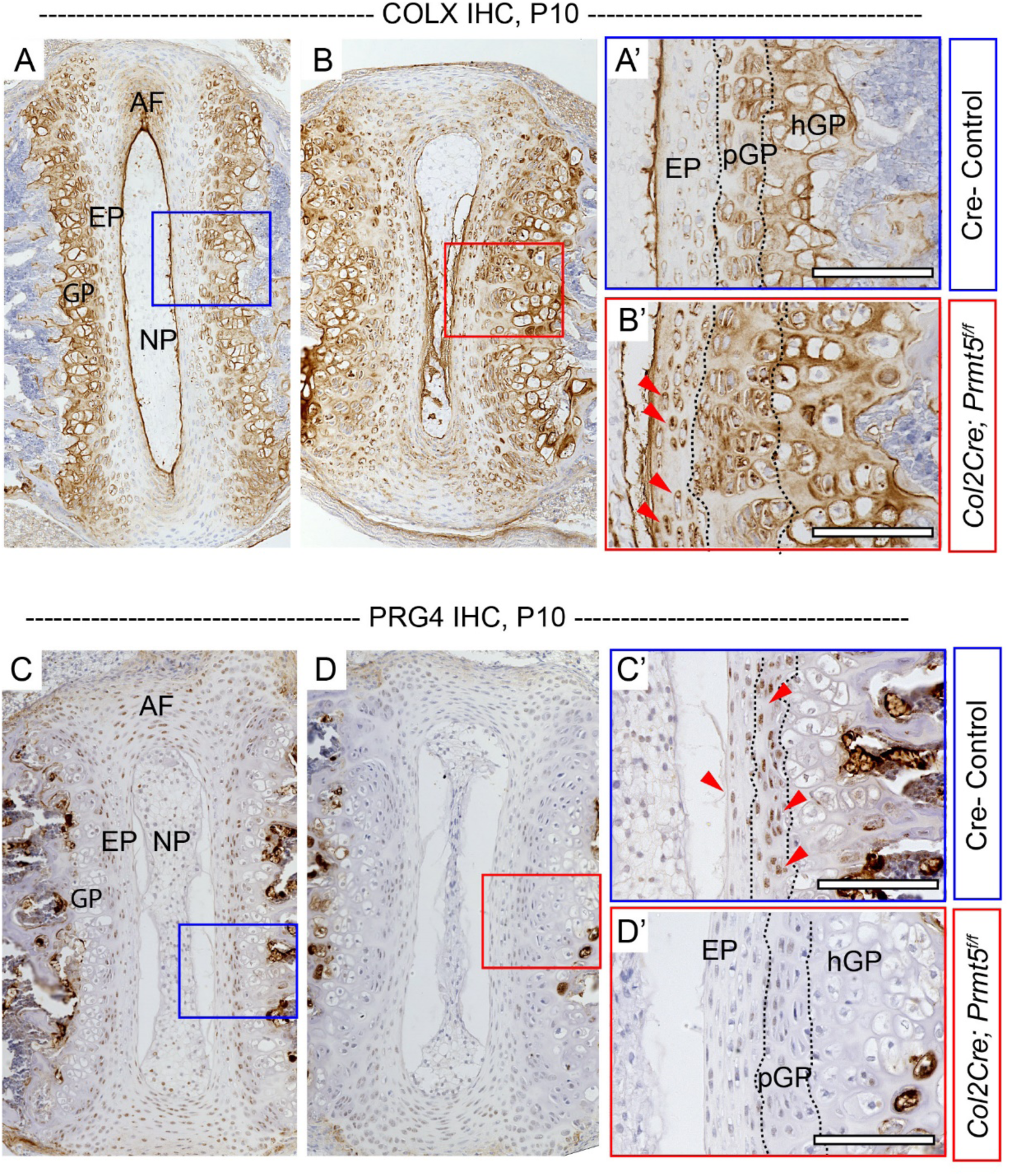
Loss of *Prmt5* in osteochondral progenitors results in induced expression of COLX and depleted expression of PRG4 in the intervertebral disc. **(A-D)** Immunohistochemistry (IHC) analysis of analysis of type X collagen (COLX) (A, B) and proteoglycan 4 (PRG4) (C, D) on spine sections of Cre- control (A and C) or *Col2Cre;Prmt5^f/f^* mutant (B and D) mice at P10. Images with higher magnification were shown in (A’-D’). Mutant mice demonstrated increased COLX signal in the endplate (red arrowheads; B’) and diminished PRG4 signal in endplate and growth plate (D’), which was consistently observed in the discs in Cre -control mice (red arrowheads; C’). (*n*=3 mice for each group.) Scale bars: 100μm in (A-D’). *AF: annulus fibrosus, EP: endplate, GP: growth plate, NP: nucleus pulposus, pGP: proliferative growth plate, hGP: hypertrophic growth plate*.

### PRMT5 regulates the expression of critical factors of chondrocyte differentiation during endochondral ossification in the spine

We next set out to determine the mechanisms by which PRMT5 regulates terminal hypertrophic differentiation. RUNX2 is a critical driver of terminal differentiation of hypertrophic chondrocytes and osteoblast differentiation in mouse (Long and Ornitz, 2013; Takarada et al., 2013), and acts in part through the activation of *Mmp13* (Inada et al., 1999; Komori, 2010a; Komori, 2018). The expression of RUNX2 was markedly diminished in *Col2Cre;Prmt5^f/f^* mutant IVD at P10 (Fig. *4B, B’*), while RUNX2 expression in control mice remained robust throughout the IVD (Fig. *4A, A’*). In contrast, we did not observe alterations of RUNX2 expression earlier in development in *Col2Cre;Prmt5^f/f^* mutant IVDs at P1 (Supplemental Fig. *5A, B*). Next, we performed fluorescent *in situ* hybridization (FISH) using an *Mmp13* specific riboprobe on medial sectioned IVDs. At P10, there was an obvious reduction of *Mmp13* expression in the growth plate and trabecular bone of the *Col2Cre;Prmt5^f/f^* mutants (Fig. *4D-D”*) compared with typical robust expression pattern in the hypertrophic zone in control IVDs (Fig. *4C-C”*). *Col2Cre;Prmt5^f/f^* mutant mice also had an obvious decrease in *Mmp13* expression in the presumptive center of ossification and vertebral growth plate at P1 (Supplementary Fig. *5D-D”*), compared with a robust *Mmp13* signal in ossification center and hypertrophic growth plate in control mice (Supplementary Fig. *5C-C”*). Interestingly, we observed ectopic *Mmp13* signal within the nucleus pulposus and the endplate in mutant IVDs at P1 (Supplemental Fig. *5D-D”*), which was not observed in control mice. The consistent reduction of RUNX2 and - its downstream effector - *Mmp13* in the IVD and vertebral growth plate of *Col2Cre;Prmt5^f/f^* mutant mice supports a mechanistic role for PRMT5 in the regulation of hypertrophic chondrocytes differentiation and endochondral ossification during perinatal development of the spine.

**Figure 4.**
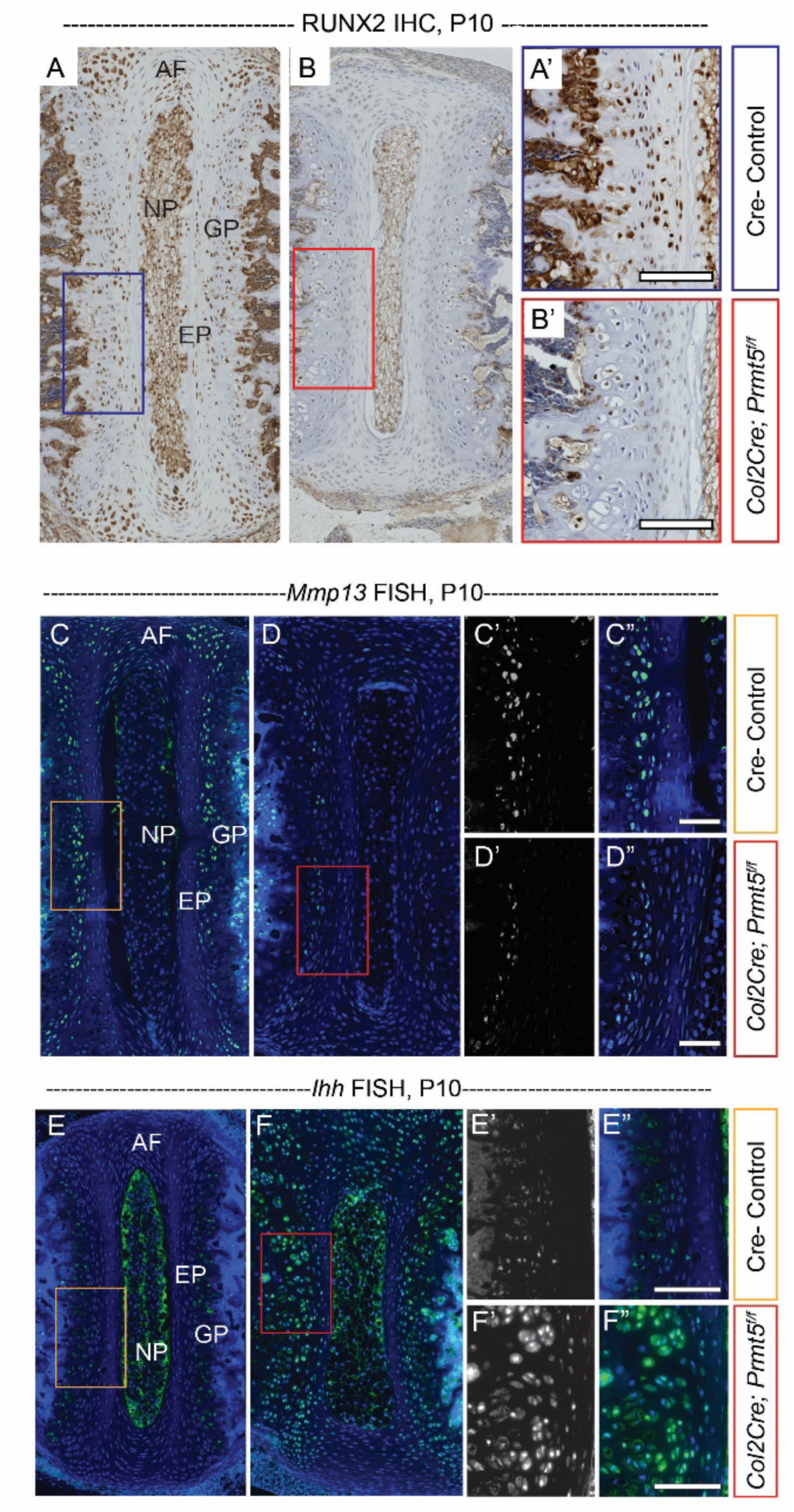
PRMT5 regulates the expression of RUNX2, *Mmp13*, and *Ihh*. **(A, B)** Immunohistochemistry (IHC) analysis of RUNX2 on spine sections of Cre - control (A) or *Col2Cre;Prmt5^f/f^* mutant (B) mice at P10 demonstrating reduced RUNX2 expression in the endplate and growth plate of mutant mice. Images with higher magnification were shown in (A’, B’). **(C, D)** Fluorescent *in situ* hybridization (FISH) analysis of *Mmp13* on spine sections of Cre- control (C) or *Col2Cre;Prmt5^f/f^* mutant (D) mice at P10. Reduced *Mmp13* signal was detected in the growth plate of the mutant mice. (C’, D’) are *Mmp13* fluorescent *in situ* channels, and (C, C”, D, D”) are merged channels. Images with higher magnification were shown in (C’, C” and D’, D”). **(E, F)** FISH analysis of *Ihh* on spine sections of Cre- control (E) or *Col2Cre;Prmt5^f/f^* mutant (F) mice at P10. Induced expression of *Ihh* was detected in the growth plate of the mutant mice. (E’, F’) are *Ihh* fluorescent *in situ* channels, and (E, E”, F, F”) are merged channels. Images with higher magnification were shown in (E’, E” and F’, F”). (*n*=3 for each group.) Scale bars: 100μm. *AF: annulus fibrosus, NP: nucleus purposes, EP: endplate, GP: growth plate*.

To better understand the regulatory mechanisms of PRMT5 during endochondral bone formation, we next assayed Indian hedgehog (*Ihh*) expression in both control and *Col2Cre;Prmt5^f/f^* mutant mice. *Ihh* signaling maintains chondrocyte proliferation and negatively regulates chondrocyte hypertrophy by regulation of PTHrP expression (Long and Ornitz, 2013). On the other hand, *Ihh* acts independently of PTHrP to promote chondrocyte hypertrophy, and induces COLX expression through RUNX2/Smad interactions (Amano et al., 2014; Mak et al., 2008). FISH analysis with an *Ihh* specific riboprobe shows comparable expression pattern in both control and *Col2Cre;Prmt5^f/f^* mutant mice at P1, confirming that loss of PRMT5 does not affect early hypertrophic differentiation (Supplemental Fig. *5E, F*). However, at P10 we observed a dramatic increase of *Ihh* expression in the mutant IVD and growth plate (Fig. *4F-F”*), including abnormal expression of *Ihh* in the proliferative growth plate, as well as in the majority of cells of the endplate and annulus fibrosus (Fig. *4F-F”*). Our results demonstrate that PRMT5 regulates terminal hypertrophic differentiation through positive regulation of RUNX2 and *Mmp13* expression and by negative regulation of *Ihh* expression during perinatal development, resulting in an obvious delay of terminal hypertrophic chondrocyte differentiation. We suggest this is in part due to the loss of RUNX2 and *Mmp13* expression which promote terminal hypertrophic chondrocyte differentiation as well as, due to increased, ectopic *Ihh* expression may act to counter hypertrophic differentiation. Taken together, our results suggest that misregulation of several critical regulators of hypertrophic chondrocyte differentiation leads to attenuated terminal hypertrophic differentiation in vertebral growth plate, changes in gene expression in the IVD, and ultimately leads to impaired endochondral ossification during perinatal spine development (see discussion).

### PRMT5 regulates *Bmp4* expression in the spine

Loss of *Prmt5* in limb bud and lung tissue induces ectopic, elevated *Bmp4* expression which is accompanied by increased apoptosis in these tissues (Li et al., 2018; Norrie et al., 2016). Consistent with previous observations in *Prmt5*-deficient limb (Norrie et al., 2016) and lung epithelium (Li et al., 2018) in mouse, we also observed increased ectopic *Bmp4* signal in the growth plate of *Col2Cre;Prmt5^f/f^* mutant IVDs at P10 that was not present in controls (Supplemental Fig. *6B, B’*). We did not observe upregulation of *Bmp4* in the IVD at P1 mice in either genotype (Supplemental Fig. *6C-D’*), suggesting that this increased expression accumulates during perinatal development. We also detected a mild increase in cell death in the IVDs of *Col2Cre;Prmt5^f/f^* mutant mice in the annulus fibrosus, endplate, and growth plate regions at both P1 and P10 (Supplemental Fig. *7B, D, E, F*). Taken together, these data demonstrate that as in other tissues and organs PRMT5 is required for regulation of *Bmp4* and apoptosis in the cartilaginous tissues of the IVD and growth plate.

### PRMT5 is not required in the mature osteoblast lineages for spine development and stability

Giving the clear alterations in endochondral bone formation in *Col2Cre;Prmt5^f/f^* mutant mice, we next sought to determine if PRMT5 also functions in committed bone-forming lineages. To address this, we utilized an *Osteocalcin (Oc) Cre* transgenic mouse to specifically remove PRMT5 function in mature osteoblasts (Zhang et al., 2002). These conditional mutant mice were born at Mendelian ratios and were all adult viable, displaying no obvious phenotypes (*n*=7). X-ray analysis showed no scoliosis in the *OcCre;Prmt5^f/f^* mutant mice at P10 (Supplemental Fig. *8A, B*, *n*=4) or at 2 months of age (data not shown, *n*=3). Histological analysis at P10 showed no obvious alterations of spine tissues in *OcCre;Prmt5^f/f^* mutant mice (Supplemental Fig. *8C, D*). As described previously we also did not detect PRMT5 expression in the cortical or trabecular bone of the spine (Supplemental Fig. *2*). Taken together, we conclude that PRMT5 functions in chondrocyte lineages to promote endochondral ossification, and does not have an obvious role in mature osteoblast lineages for this process.

### PRMT5 has a role in homeostasis of the adult intervertebral disc

To gain insight into whether PRMT5 has a role in homeostasis of the spine in adult mice, we assayed the expression of PRMT5 in wild-type IVD by IHC at 6-weeks and 4-months-of-age. We observed low level of PRMT5 expression in a group of cells bordering the ligament insertion and perichondrium region of the IVD at both time points (Supplemental Fig. *9*).

Given indication of continued PRMT5 in adult IVD, we decided to assay the role of PRMT5 in spinal homeostasis. To specifically remove PRMT5 function in cartilaginous tissues of the spine in adult mouse, we cross *Prmt5^f/f^* mice with an *Acan enhancer-driven, Tetracycline-inducible Cre* (*ATC*) transgenic mouse which targets committed chondrocyte lineages (Dy et al., 2012), and induced recombination from 4 to 8 weeks of age (Fig. *5A*). Beta-galactosidase staining in *ATC;Rosa-LacZ ^f/+^* mice revealed near complete recombination in nucleus pulposus, endplate, and annulus fibrosus of the IVD, as wells as over 50% recombination in growth plate of the at one week post injection (5 weeks-of-age) using this protocol for recombination (Fig. *5C*). We first assayed several markers of IVD using qPCR analysis of whole IVD cDNA libraries in experimental groups at 6 weeks post injection (2.5 months-of-age) (Fig. *5A*). We found that consistent with the observations in *Col2Cre;Prmt5^f/f^* mutant mice, there was a ∼2-fold reduction in *Prmt5* expression, as well as ∼2-fold reduction in the expression of *Mmp13* and *Prg4*, respectively. In contrast, the expression of *Sox9*, *Col2a1*, *Acan,* and *Col10a1* were not strongly affected at this time point (Fig. *5D*). By the age of 4 months (3 months post induction), there were no overt signs of degenerative histopathology in *ATC;Prmt5^f/f^* mutant IVD (Fig. *5F*), with the exception of a minor increase of acellular clefts at the midline of the endplate (yellow arrow; Fig. *5F*), which we occasionally observe in wild type IVD as well (*p*=0.047; two-tailed student *t* test; *n*=3 mice for each group). However, we observed increased, ectopic COLX expression in the endplate and growth plate of *ATC;Prmt5^f/f^* mutant IVDs (Fig. *5H, H’*). Taken together, our data shows that loss of PRMT5 in adult IVD results in reduced expression of normal extracellular matrix component *Prg4*, coupled with increased expression of hypertrophic marker COLX and decreased expression of *Mmp13*, consistent with a continuous role for PRMT5 in regulating hypertrophic chondrocyte turnover and homeostasis of the IVD in adulthood. We did not observe scoliosis in *ATC;Prmt5^f/f^* mutant mice when induced from either 2-weeks-of-age or 4-weeks-of-age by dorsal X-ray imaging (Supplemental Fig. *10* and data not shown). Taken together, our study demonstrates a novel role for PRMT5 in cartilaginous lineages for the regulation of adult IVD homeostasis, but has a limited temporal role during perinatal development for the regulation of spine stability.

**Figure 5.**
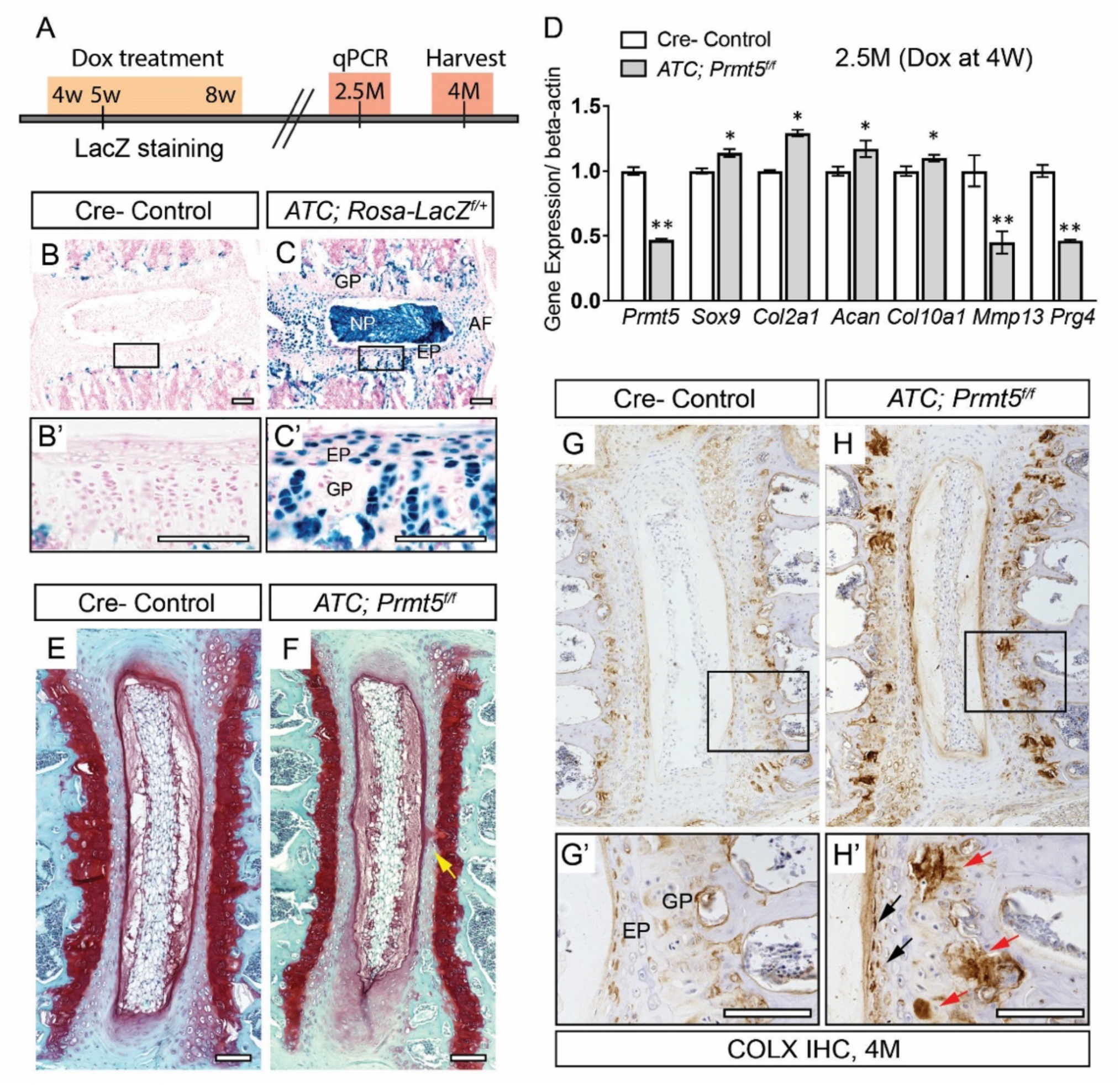
Postnatal ablation of *Prmt5* in the IVD induces the onset of markers of disc degeneration. **(A)** Schematic of protocol for postnatal recombination of the floxed PRMT5 allele used for these experiments. **(B, C)** Beta-galactosidase staining on spine sections of Cre - control (B) or *ATC;Rosa-LacZ ^f/+^* (C) mice at 5-weeks of age shows that effective recombination is achieved within the IVD at 1-week post Dox induction. Images with higher magnification were indicated with black boxes and shown in B’, C’. **(D)** qPCR analysis of IVD tissue harvested at 2.5-months of age demonstrates effective reduction in *Prmt5* as well as *Mmp13* and *Prg4*. Bars represent mean ± SD. (*n*=3 for each group.) “*” indicates p<0.05, “**” indicates p<0.01, two-tailed student *t*-test. **(E, F)** Safranin O/ Fast Green staining on medial sectioned IVD of Cre -control (E) or *ATC; Prmt5^f/f^* mutant (F) mice harvested at 4-months of age after Dox treatment. No obvious histopathology was observed except for the occurrence of minor clefts of the endplate and growth plate in mutant mice (yellow arrow; F). (G, H) Immunohistochemistry of COLX protein on medial sectioned IVD from Cre - control (G) or *ATC; Prmt5^f/f^* mutant (H) mice at 4-months of age. Images with higher magnification were indicated with black boxes and shown in G’, H’. Increased and ectopic COLX signal was observed in endplate (H’, black arrows) and growth plate (H’, red arrows) of the mutant IVD. (*n*=3 for each group.) Scale bars: 100μm. *AF: annulus fibrosus, EP: endplate, GP: growth plate, NP: nucleus pulposus*.

## DISCUSSION

The formation of the vertebral bodies, like the long bones, ossify through a process of endochondral ossification where hypertrophic chondrocytes are replaced by and in some cases contribute to bone-forming osteoblast lineages (Aghajanian and Mohan, 2018). Our findings indicate that PRMT5 has an important role in the process of terminal differentiation of hypertrophic chondrocytes and endochondral bone formation, in part by positive regulation *Mmp13* and RUNX2 expression and negative regulation of *Ihh* expression. We observed that loss of PRMT5 function in osteochondral progenitors results in the expansion of hypertrophic chondrocytes in the vertebral growth plate. However, ossification of the vertebrae is severely impaired during perinatal development, mimicking an *Mmp13* loss-of-function phenotype (Inada et al., 2004). Finally, we show that PRMT5 regulation of perinatal endochondral bone formation is a potential mechanism underlying infantile IS. Taken together, these results establish PRMT5 as a fundamental regulator of terminal hypertrophic chondrocyte differentiation and endochondral bone formation, as well as a critical regulator for perinatal spine integrity.

### Regulation of Terminal Chondrocyte Differentiation and Endochondral Bone Formation by PRMT5

One of the most important regulators of hypertrophic chondrocyte differentiation and endochondral bone formation is RUNX2 (Komori, 2010b; Komori, 2018). *Runx2*-deficient mice exhibit a complete loss of mature osteoblasts and endochondral bone formation (Chen et al., 2014; Ducy et al., 1997; Takarada et al., 2013). RUNX2 has been shown to directly regulate *Col10a1*, *Mmp13* and *Vegfa* expression (Hess et al., 2001; Li et al., 2011; Takahashi et al., 2017; Zelzer et al., 2001), and therefore plays a crucial role in driving chondrocyte differentiation and endochondral ossification (Fig. *6A*). *Mmp13*, a direct target of RUNX2 (Fig. *6A*), is also required for these processes as mice lacking *Mmp13* display expanded hypertrophic zone and delay in endochondral ossification in the long bone (Inada et al., 2004; Stickens et al., 2004). Here, we found that *Col2Cre;Prmt5^f/f^* mutant mice displayed an obvious reduction in RUNX2 and *Mmp13* expression in proliferative and hypertrophic chondrocytes of the perinatal growth plate (Fig. *4* and Supplemental Fig. *5*). Meanwhile, the *Col2Cre;Prmt5^f/f^* mutant mice showed an expansion of the hypertrophic growth plate with ectopic, expanded COLX expression. These results indicate that loss of PRMT5 in osteochondral progenitors allows hypertrophic differentiation to commence; however, the process of removing COLX positive hypertrophic cells is severely impaired due to loss of RUNX2/*Mmp13* expression (Fig. *6B*). How hypertrophic chondrocyte differentiation is able to proceed, albeit in a severely delayed manner, without these factors is still under investigation. It is possible that undetectable levels of these proteins are being made or that other RUNX-family members or co-factors that are important for driving terminal differentiation, such as RUNX3, MEF2C, or FOXA2 as indicated in long bone (Arnold et al., 2007; Tan et al., 2018; Yoshida et al., 2004), are able to act in a semi-redundant manner for this process.

**Figure 6.**
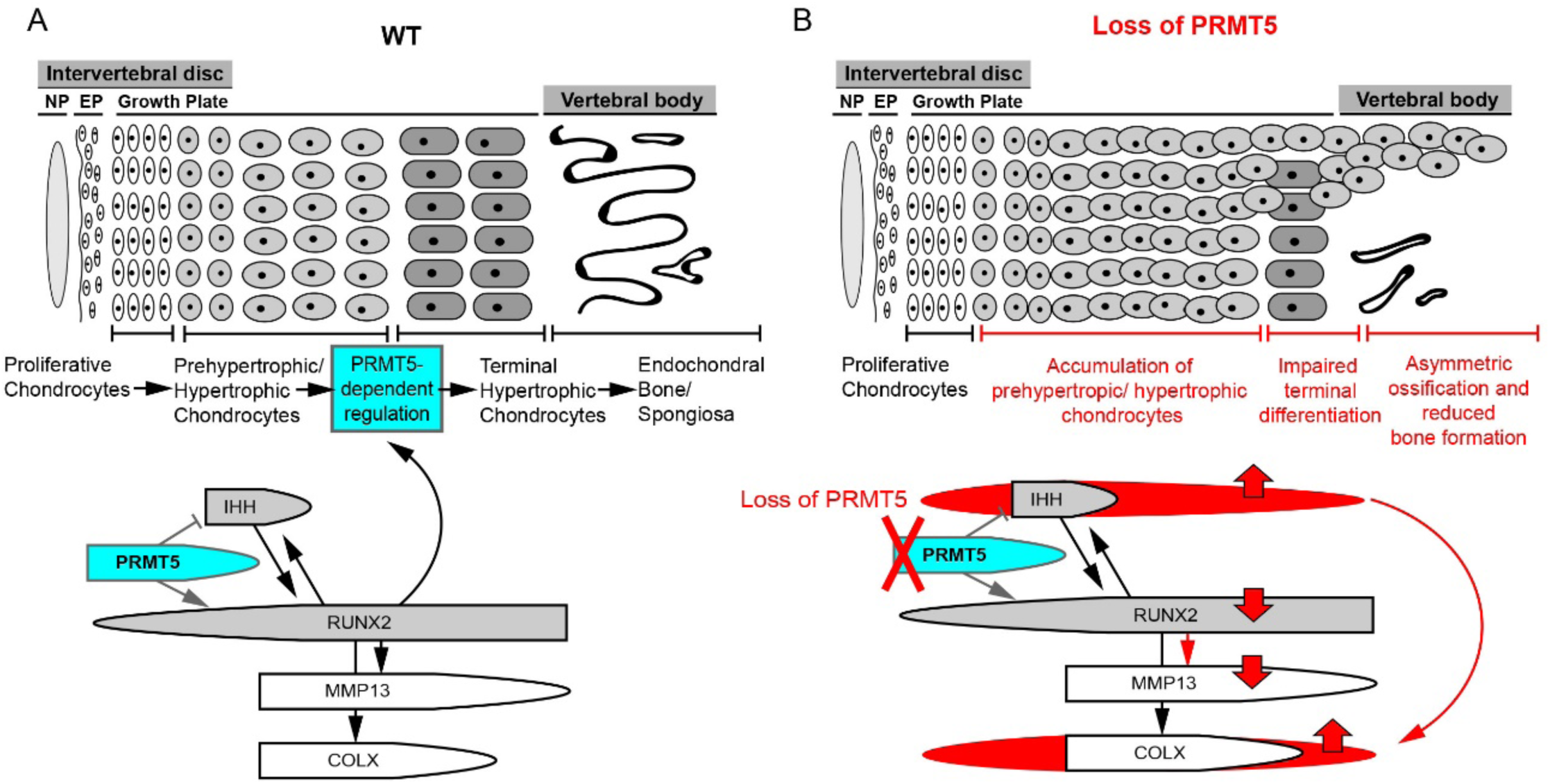
A model of PRMT5 in regulating terminal differentiation of hypertrophic chondrocytes in the perinatal growth plate of the spine. **(A)** PRMT5 expressed in proliferative and prehypertrophic chondrocytes functions to regulate chondrocyte hypertrophic differentiation by controlling RUNX2 and IHH signaling. PRMT5 is required for maintaining postnatal expression of RUNX2, which is a key regulator for terminal differentiation of hypertrophic chondrocytes, and a positive regulator of MMP13 expression. PRMT5 also negatively regulate the expression of *Ihh*. (B) *Prmt5* deletion in cartilaginous tissues of the spine leads to expanded *Ihh* expression in perinatal growth plate and promotes the expression/accumulation of COLX. It also results in reduced RUNX2 and *Mmp13* expression, which inhibits the terminal differentiation of the hypertrophic chondrocytes. These two processes synergistically leads to persistent hypertrophic growth plate and reduced bone formation in perinatal mouse vertebra. *NP: nucleus pulposus, EP: endplate*.

Another important regulator of hypertrophic chondrocyte differentiation is IHH. *Ihh* signaling plays critical role in both chondrocyte proliferation and hypertrophic differentiation. It can directly promote proliferation and inhibit premature hypertrophy of the proliferative chondrocyte via IHH/PTHrP negative feedback loop (Long and Ornitz, 2013). On the other hand, it can promote hypertrophy and induce COLX expression in a PTHrP independent manner (Amano et al., 2014; Mak et al., 2008). We observed that *Col2Cre;Prmt5^f/f^* mutant mice displayed ectopic expansion of *Ihh* expression throughout the hypertrophic growth plate, proliferative growth plate, and some cells within the endplate of IVD at perinatal (P10) stage but not neonatal stage (P1) (Fig. *4F* and Supplemental Fig. *5F*), which overlaps with the regions of ectopic, expanded COLX expression (Fig. *2B* and Fig. *6B*). These results indicate that loss of PRMT5 in osteochondral progenitors allows chondrocyte proliferation and hypertrophic differentiation, however it fails to shut down/restrict *Ihh* expression during perinatal development, which may contribute to increased production and accumulation of COLX positive cells in the growth plate. We speculate that the misregulation of *Mmp13* and *Ihh* in *Col2Cre;Prmt5^f/f^* mutant mouse spines act synergistically to promote the accumulation of hypertrophic chondrocytes in the growth plate and inhibit their terminal differentiation, which leads to reduced endochondral bone formation during perinatal skeletogenesis.

In addition, PRMT5 has been shown to regulate *Bmp4* expression in several contexts. For example, loss of *Prmt5* in mouse mesenchymal progenitor cells led to upregulated *Bmp4* in mouse limb (Norrie et al., 2016) and PRMT5 directly associated with chromatin of *Bmp4* to suppress its transcription during lung branching morphogenesis (Li et al., 2018). In agreement, *Col2Cre;Prmt5^f/f^* mutant mice also display ectopic elevation of *Bmp4* expression in the hypertrophic growth plate (Supplemental Fig. *6*). Interestingly, *in vitro* studies have shown that *Bmp4* can stimulate chondrocyte hypertrophy and induce COLX expression (Clark et al., 2009; Hatakeyama et al., 2004; Minina et al., 2001; Steinert et al., 2009). However, whether PRMT5-dependent regulation of *Bmp4* expression is related to defective hypertrophic chondrocyte differentiation remains to be determined.

Interestingly, the formation of the cartilaginous templates and some ossification occurs normally in these conditional *Col2Cre;Prmt5^f/f^* mutant mice (Fig. *1C*) despite *Prmt5* deletion in osteochondral progenitor lineages. This is in contrast to the more severe loss of cartilage template in *Prx1Cre;Prmt5^f/f^,* resulting in a dramatic reduction of the mouse forelimb (Norrie et al., 2016). Taken together, these findings support a model of distinct mechanistic roles PRMT5 function (i) for initiation of early chondrocyte progenitors and (ii) for regulation of hypertrophic chondrocyte differentiation.

### PRMT5 regulates homeostasis of the adult IVD

We found that postnatal loss of PRMT5 in cartilaginous tissues of the IVD resulted in alterations of the normal gene expression including reduced *Prg4* expression as well as increased COLX expression, which are both markers of early-onset degenerative disc. These findings demonstrate that PRMT5 has an important role in the homeostasis of cartilaginous tissues of the IVD. It will be important to determine if the ablation of PRMT5 in the IVD of mature adult mice can generate susceptibility to the onset of pathogenic changes of the IVD and spine due to trauma or aging in mice. While the IVD has been considered an organ with little or no regenerative capacity, several studies have recently identified the presence of cells expressing stem/progenitor markers in this tissue (Blanco et al., 2010; Henriksson and Brisby, 2013; Risbud et al., 2007). Interestingly, a study using BrdU labeling identified label-retaining cells in the annulus fibrosus located in a region bordering the ligament insertion and the perichondrium region (Henriksson et al., 2009). This region overlaps with the expression of PRMT5 in the adult IVD in mouse (Supplemental Fig. *9*), and suggests a possible role for PRMT5 in maintaining adult progenitor/stem cell pools which are critical for IVD homeostasis. Additional labeling studies will be necessary to test this model.

### Asymmetrical properties of the spine result in scoliosis

The molecular genetics and underlying pathology of IS are largely unknown, more so for infantile onset forms of the disease. Asymmetries of the vertebral body growth and of the mechanical properties of the bony prominences, cartilaginous joints of the vertebral bodies, and musculature attachments of the spinal column have long been hypothesized to underlie the formation of idiopathic scoliosis (Liu and Gray, 2018). Our results suggest that the onset of infantile IS in *Col2Cre;Prmt5^f/f^* mutant mice is the result of obvious defects in endochondral ossification of the spine. We suggest a model where asymmetrical defects of ossification of the vertebrae result in anisotropic mechanical properties of the spinal column causing vertebral rotation and scoliosis of the spine during perinatal development. This model is supported by our observation of a complete lack of scoliosis in conditional *ATC;Prmt5^f/f^* mutant mice recombined at 2 or 4-weeks-of-age, which also display alterations in terminal chondrocyte differentiation, yet have a spinal column that has already undergone substantial ossification. This suggests that infantile IS may be the result in subtle defects and delays in endochondral ossification during perinatal development. Our current studies suggest that the cartilaginous tissues play a prominent role in the pathogenesis of infantile IS. In agreement, genetic defects in osteochondral progenitors of the spine inducing IS in mice also occurs in other genetic mouse models displaying IS (Liu and Gray, 2018), including conditional mutant mice of the *Gpr126*/*Adgrg6* (Karner et al., 2015), *Sox9* (Henry et al., 2012), and *Gdf5/6* (Settle et al., 2003) genes. These findings underscore the importance of proper cartilage maturation, including endochondral ossification, for maintaining spine stability during maturation and development of the spine. Taken together, our data for the first time demonstrates that PRMT5 is a critical factor for perinatal spine stability via regulation of hypertrophic differentiation pathways. Our study further demonstrates that PRMT5 may have an important function in IVD progenitor/stem cells for maintenance of IVD homeostasis. Therefore, PRMT5 may provide a key target for the development of novel therapeutics to treat infantile IS and degenerative changes to the IVDs.

## ACKNOWLEDGEMENTS

We thank Dr. Fanxin Long for sharing *Col2Cre* mice and Dr. Véronique Lefebvre for sharing *ATC* mice. We thank Dr. Matthew Hilton for providing the *Mmp13* and *Ihh in situ* probe templates. We acknowledge Dr. Courtney Karner for helpful comments on this manuscript prior to submission. Research reported in this publication was supported by the National Institute of Arthritis and Musculoskeletal and Skin Diseases of the National Institutes of Health under Award Number R01-AR072009 (R.S.G.), F32-AR073648 (Z.L.). It is also supported by NIH grant R01-HD073151 (S.V.), and a UT Austin Provost Graduate Excellence Fellowship (JR).

## Authors’ roles

Study design: ZL and RSG. Study conduct and Data collection: ZL and JR. Data analysis: ZL. Data interpretation and approving final version of manuscript: ZL, JR, SV and RSG. ZL and RSG take responsibility for the integrity of the data analysis.

---

**Supplemental Table 1.**
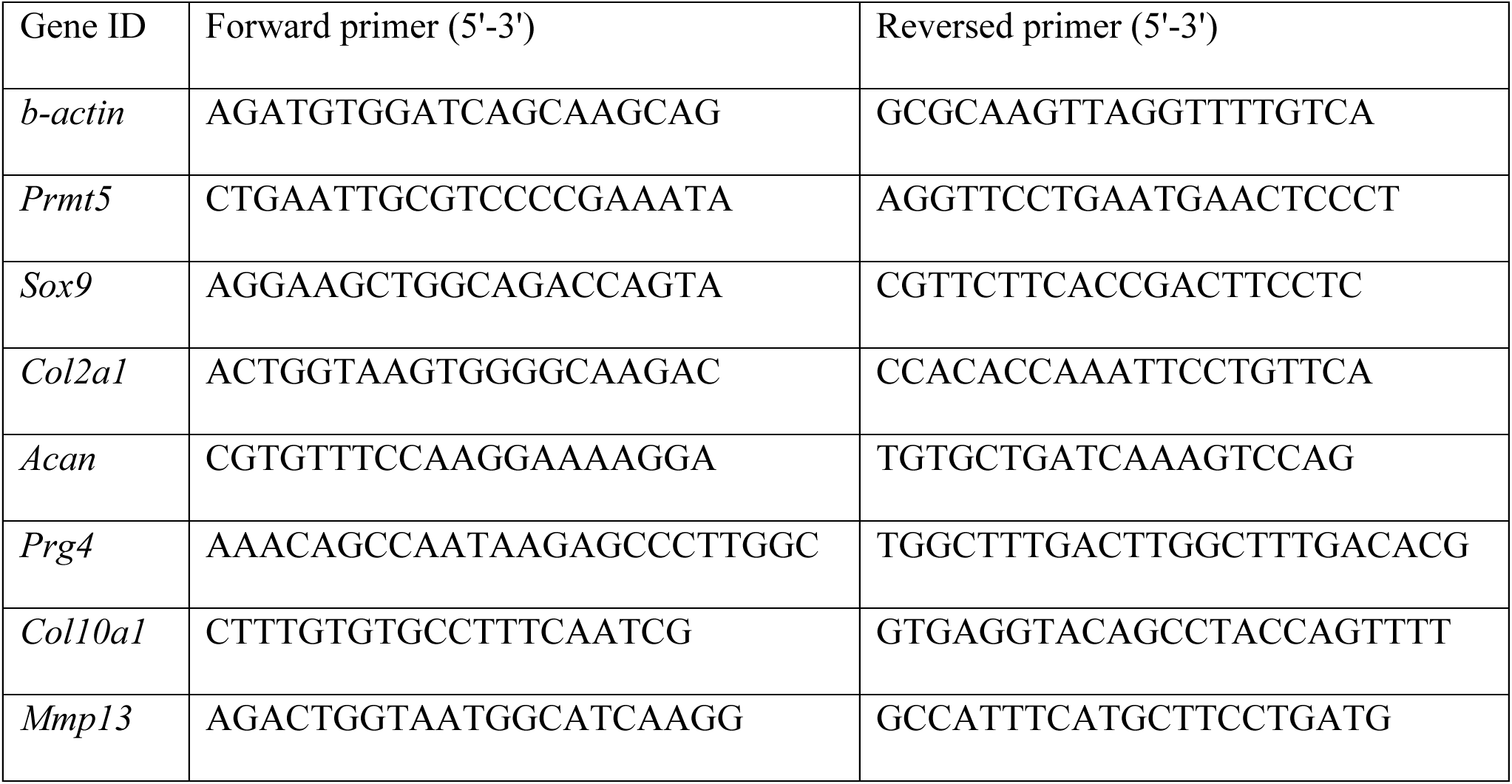
qRT-PCR primers used in this study.

**Supplemental Figure 1.**
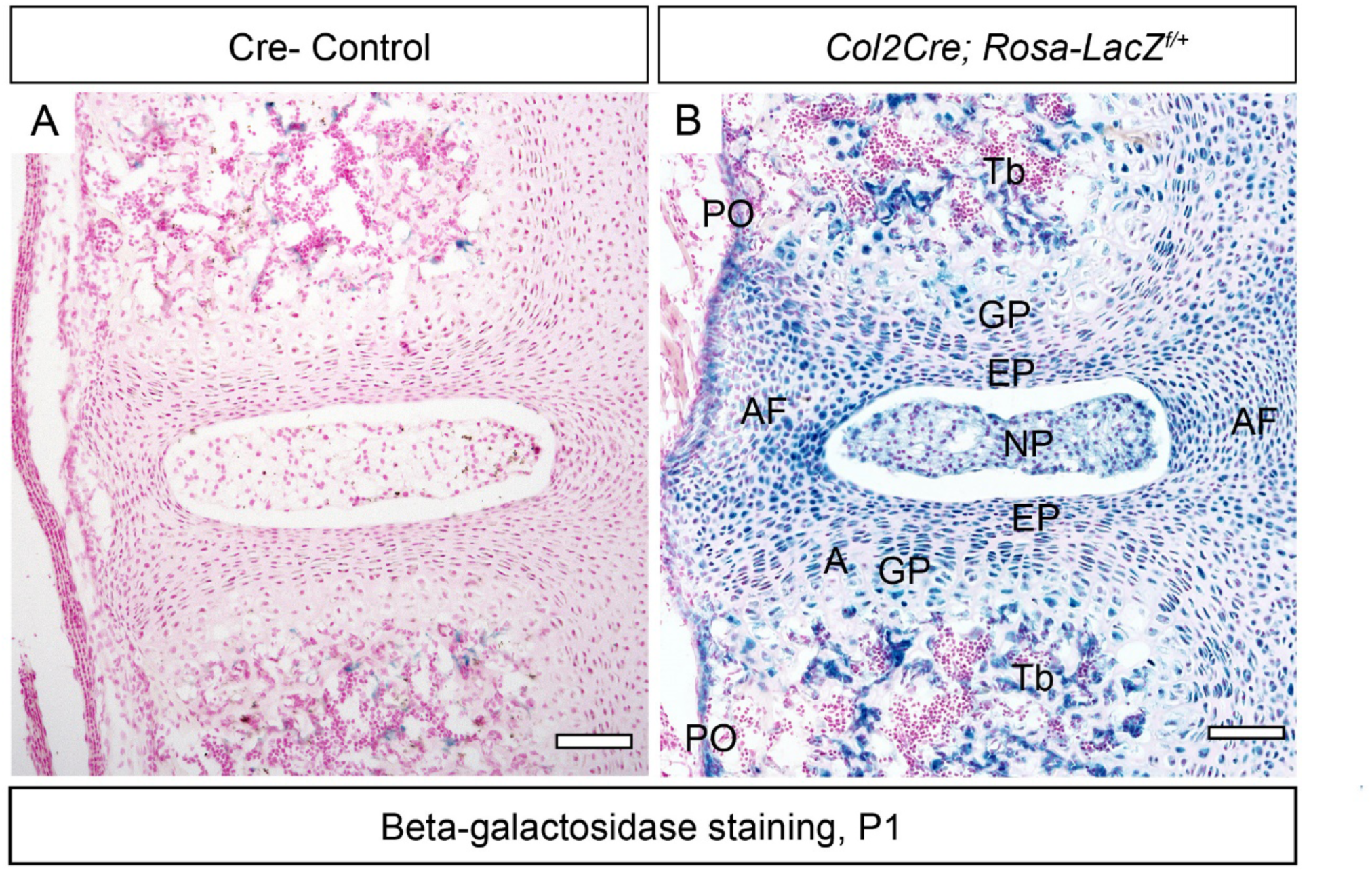
The *Col2Cre* transgene targets osteochondral progenitors. **(A, B)** Beta-galactosidase staining staining on spine sections of Cre- control (A) or *Col2Cre; Rosa-LacZ ^f/+^* mice (B) at P1. Recombination signals (blue) can be observed in the entire IVD, including nucleus purposes (NP), endplate (EP), growth plate (GP), annulus fibrosus (AF), as well as periosteum (PO) and some newly formed trabecular bone (Tb) tissues. Scale bar: 100μm.

**Supplemental Figure 2.**
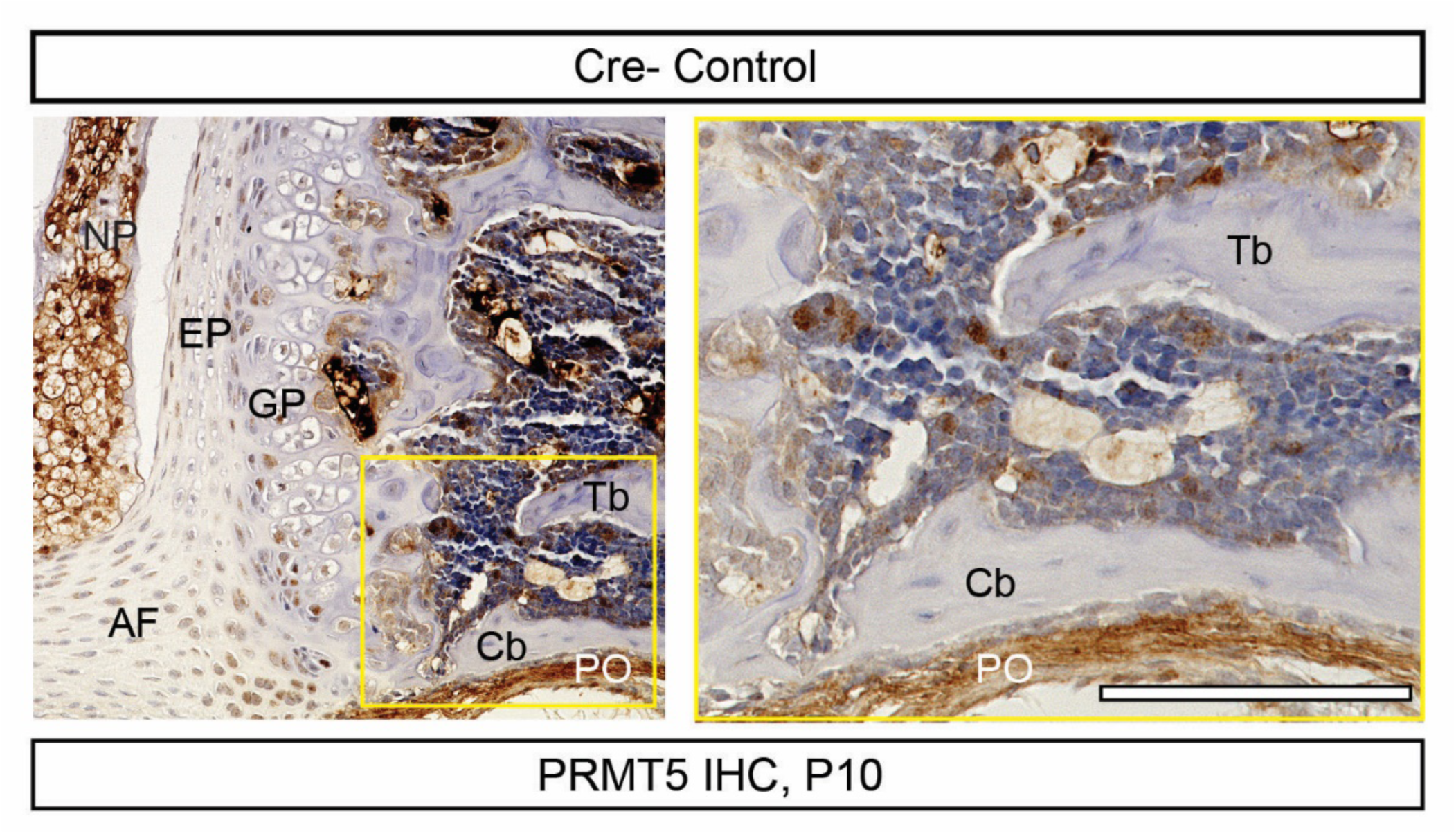
PRMT5 antibody staining on spine section. Immunohistochemistry (IHC) analyses of PRMT5 in Cre- control mice demonstrating a lack of signal in the trabecular bone or cortical bone in the control mice (*n*=3). Image with higher magnification were indicated with a yellow box. Scale bar: 100μm. *AF: annulus fibrosus, EP: endplate, GP: growth plate, NP: nucleus purposes, Tb: trabecular bone, Cb: cortical bone, PO: periosteum*.

**Supplemental Figure 3.**
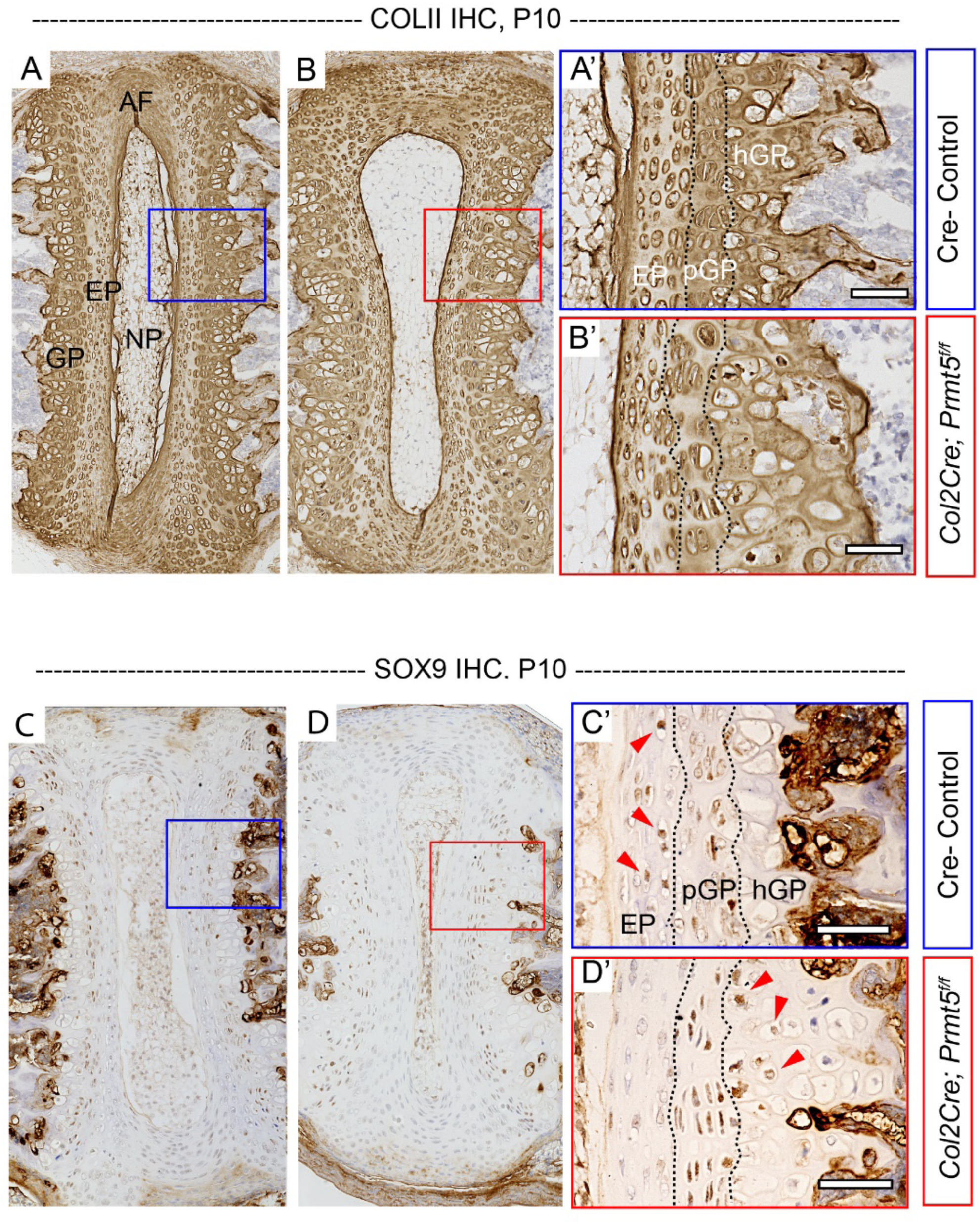
Loss of *Prmt5* in osteochondral progenitors results in mild reduction of COLII expression and abnormally expressed SOX9. **(A-D)** Immunohistochemistry (IHC) analysis of type II collagen (COLII) (A-B’) on spine sections of Cre - control (A, A’) or *Col2Cre;Prmt5^f/f^* mutant (B, B’) mice at P10 demonstrating typical COLII expression in mutant mice. **(C-D)** IHC analysis of SOX9 on spine sections of Cre (-) control (C) or *Col2Cre;Prmt5^f/f^* mutant (D) mice at P10. Images with higher magnification were shown in (A’-D’). SOX9 is normally expressed in the endplate (red arrowheads, C’) and proliferative zone of the growth plate (C’) in the control mice, but is reduced in the endplate of mutant mice (D’). Some cells in the hypertrophic zone of growth plate of mutant mice also express SOX9 (red arrowheads, D’). (*n*=3 for each group.) Scale bars: 100μm in (B’, D’). *AF: annulus fibrosus, EP: endplate, GP: growth plate, NP: nucleus pulposus, pGP: proliferative growth plate, hGP: hypertrophic growth plate*.

**Supplemental Figure 4.**
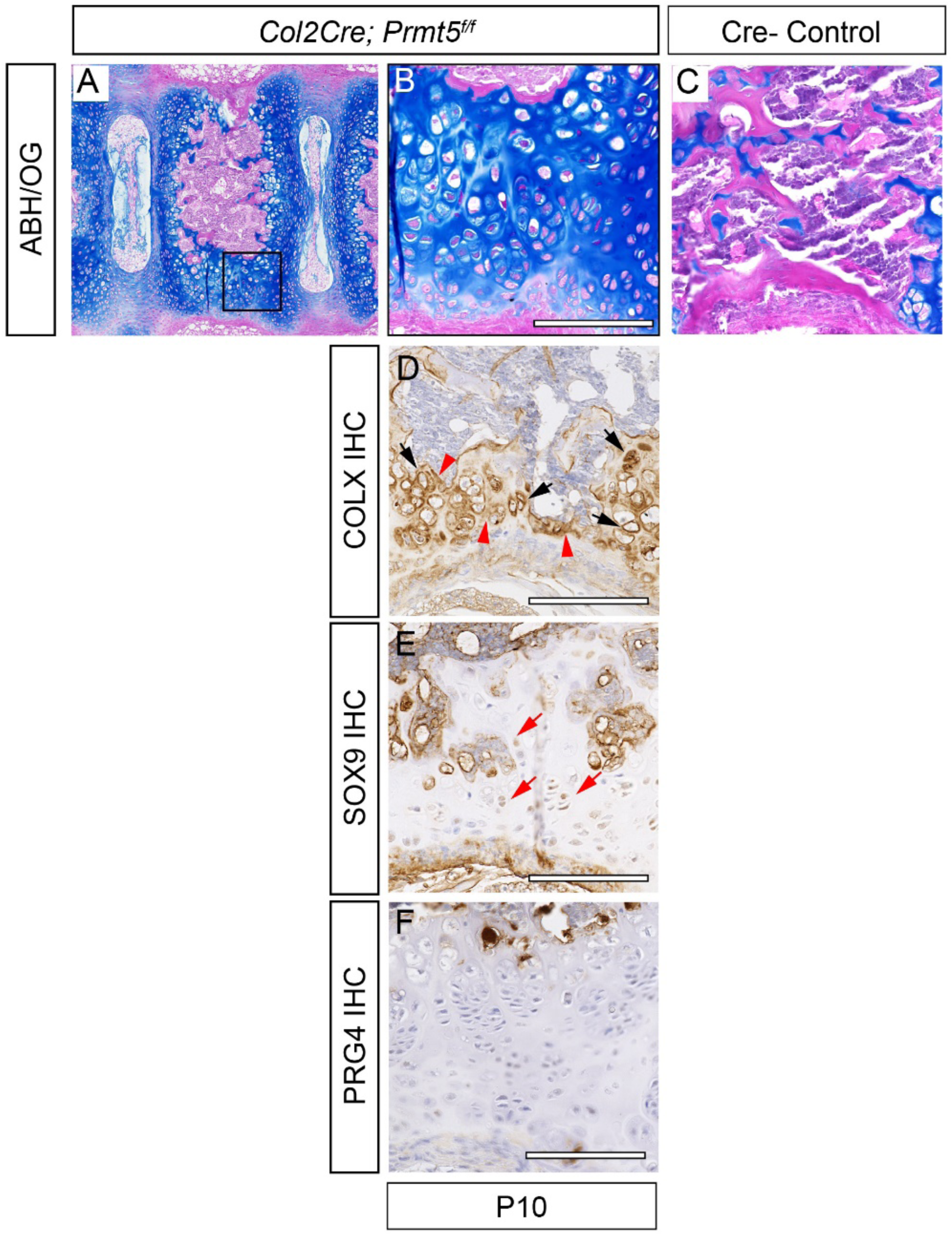
The persistent cartilage in *Col2Cre;Prmt5^f/f^* mutant mice shows abnormally expressed extracellular matrix componentts. **(A-C)** Alcian Blue Hematoxylin/ Orange G (ABH/OG) staining on spine sections of *Col2Cre;Prmt5^f/f^* mutant (A, B) and Cre- control (C) mice at P10. Persist cartilage (A, black box) was shown with higher magnification in B. There is no persist cartilage exist in Cre- control mice (C). **(D-F)** IHC analyses of COLX, SOX9, and PRG4, respectively, on adjacent sections as shown in B. The persistent cartilage highly expresses COLX in both pericellular (D, black arrows) and extracellular (D, red arrow heads) matrix, and sparkly expresses SOX9 (E, red arrows). However, the persistent cartilage does not express PRG4 (F). (*n*=3.) Scale bars: 100μm.

**Supplemental Figure 5.**
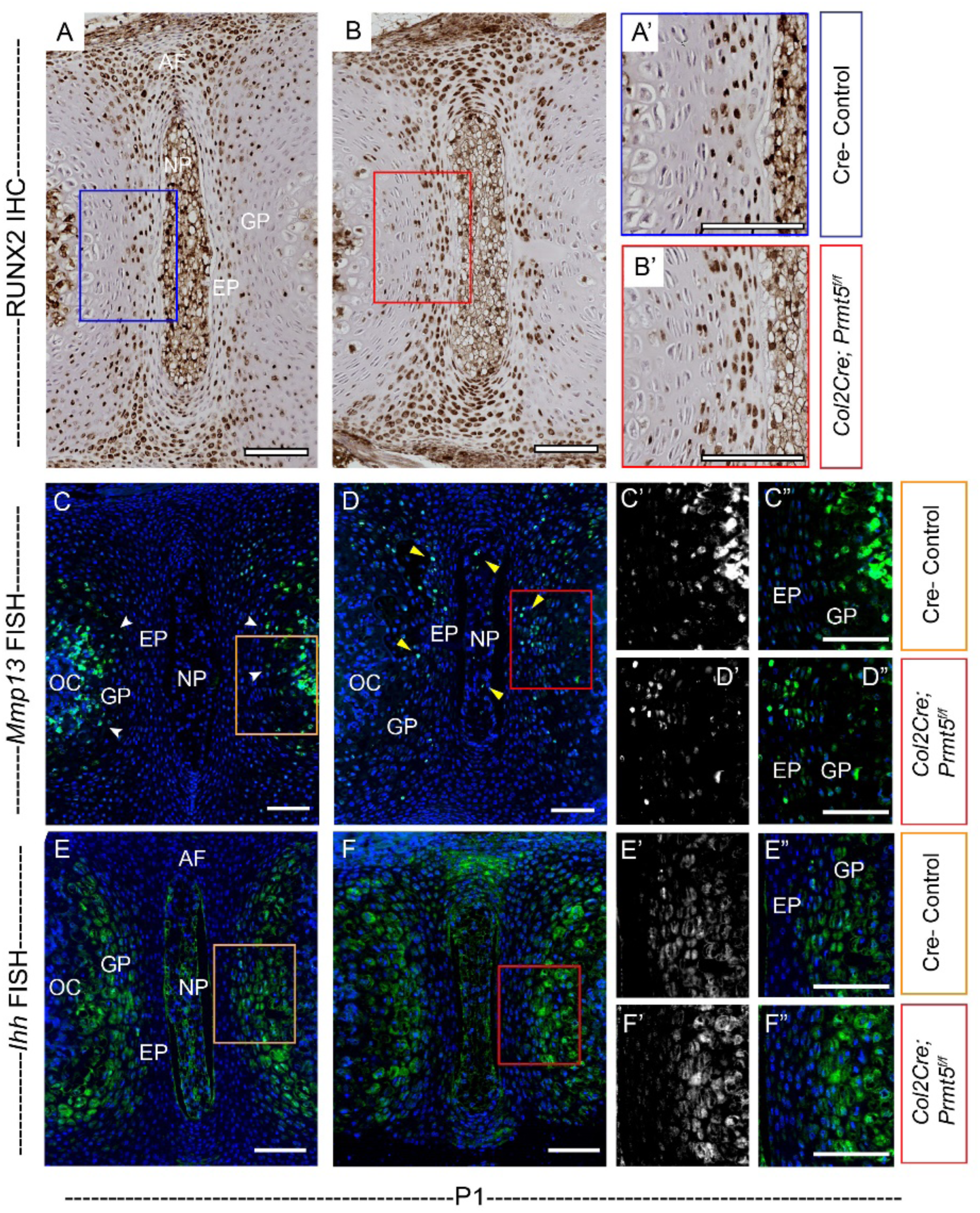
Loss of *Prmt5* in osteochondral progenitor cells results in regularly expressed RUNX2and *Ihh* but abnormally expressed *Mmp13* in newborn mice. **(A, B)** Immunohistochemistry (IHC) analysis of RUNX2 on spine sections of Cre - control (E) or *Col2Cre;Prmt5^f/f^* mutant (F) mice at P1 demonstrating typical RUNX2 expression in mutant mice. Images with higher magnification were shown in (A’, B’). **(C, D)** Fluorescent *in situ* hybridization (FISH) analysis of *Mmp13* on spine sections of Cre- control (A) or *Col2Cre;Prmt5^f/f^* mutant (B) mice at P1. Strong *Mmp13* signal was detected in the growth plate and the newly formed ossification center in the control IVD (white arrowheads; C). However, it was almost depleted in the growth plate and the OC in the mutant mice (D). On the contrary, ectopic *Mmp13* signal was observed in the endplate and the nucleus purposes in the mutant IVD (yellow arrowheads; D). (C’, D’) are *Mmp13* fluorescent *in situ* channels, and (C, C”, D, D”) are merged channels. Images with higher magnification were shown in (C’, C” and D’, D”). **(E, F)** FISH analysis of *Ihh* on spine sections of Cre- control (A) or *Col2Cre;Prmt5^f/f^* mutant (B) mice at P1. Comparable expression pattern of *Ihh* was detected in both control and mutant IVD. (E’, F’) are *Ihh* fluorescent *in situ* channels, and (E, E”, F, F”) are merged channels. Images with higher magnification were shown in (E’, E” and F’, F”). (*n*=3 for each group.) Scale bars: 100μm. *OC: ossification center, NP: nucleus purposes, EP: endplate, GP: growth plate*.

**Supplementary Fig. 6.**
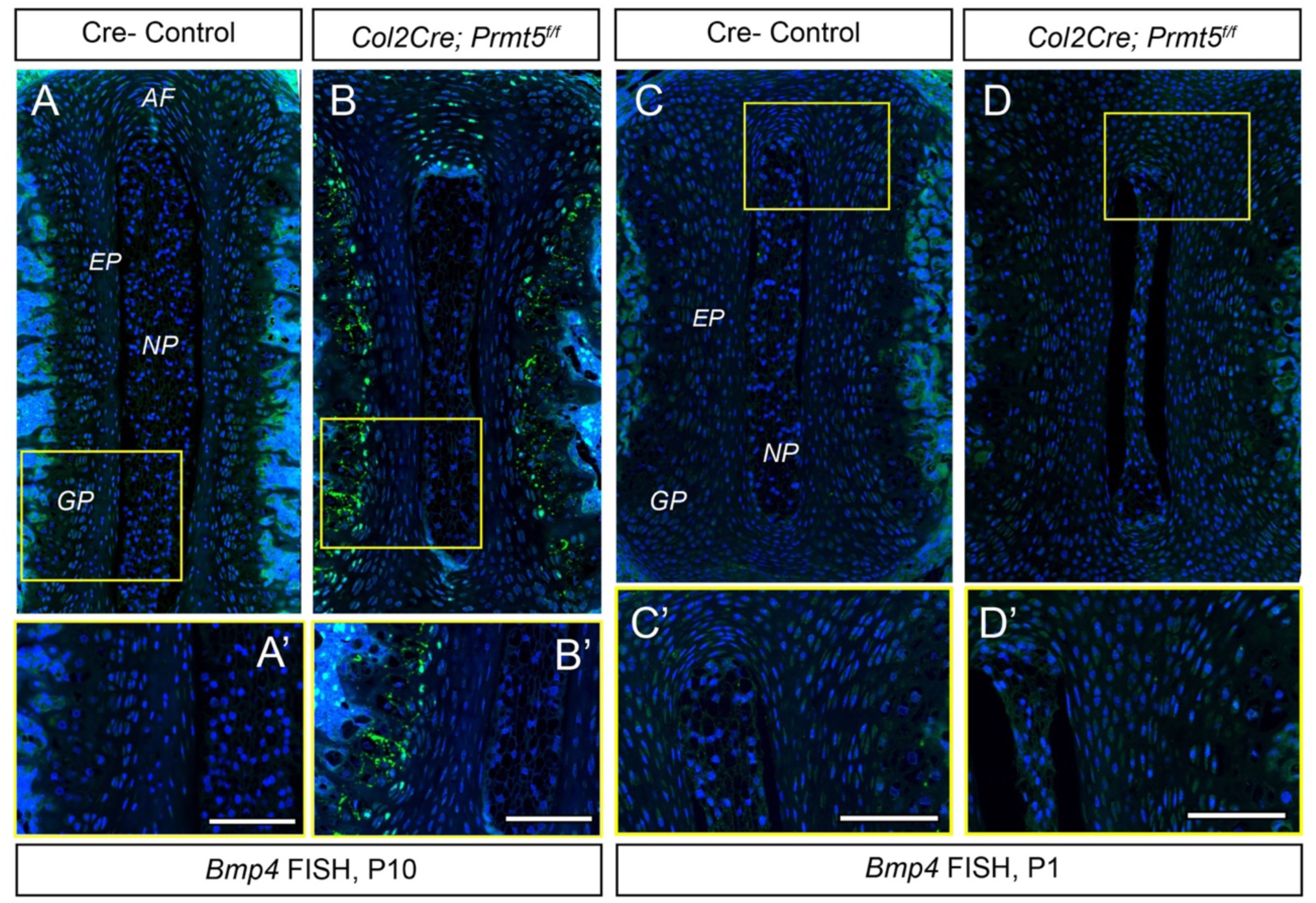
Loss of *Prmt5* in osteochondral progenitor cells induces increased, ectopic *Bmp4* expression in the intervertebral disc at P10, but not at P1. **(A-D)** Fluorescent *in situ* hybridization (FISH) analysis demonstrating *Bmp4* expression on spine sections of Cre - controls (A and C) or *Col2Cre;Prmt5^f/f^* mutant (B and D) mice. Images with higher magnification were indicated with yellow boxes and shown in A’-D’. At P10 we observe increase, ectopic *Bmp4* expression in the medial annulus fibrosus and growth plate of the IVD (B, B’) compared to an absence of expression in control mice (A, A’). However, at P1 this *Bmp4* signal was not detected in the IVD of either genotype. (*n*=3 for each group.) Scale bars: 100μm. *AF: annulus fibrosus, NP: nucleus purposes, EP: endplate, GP: growth plate*.

**Supplementary Fig. 7.**
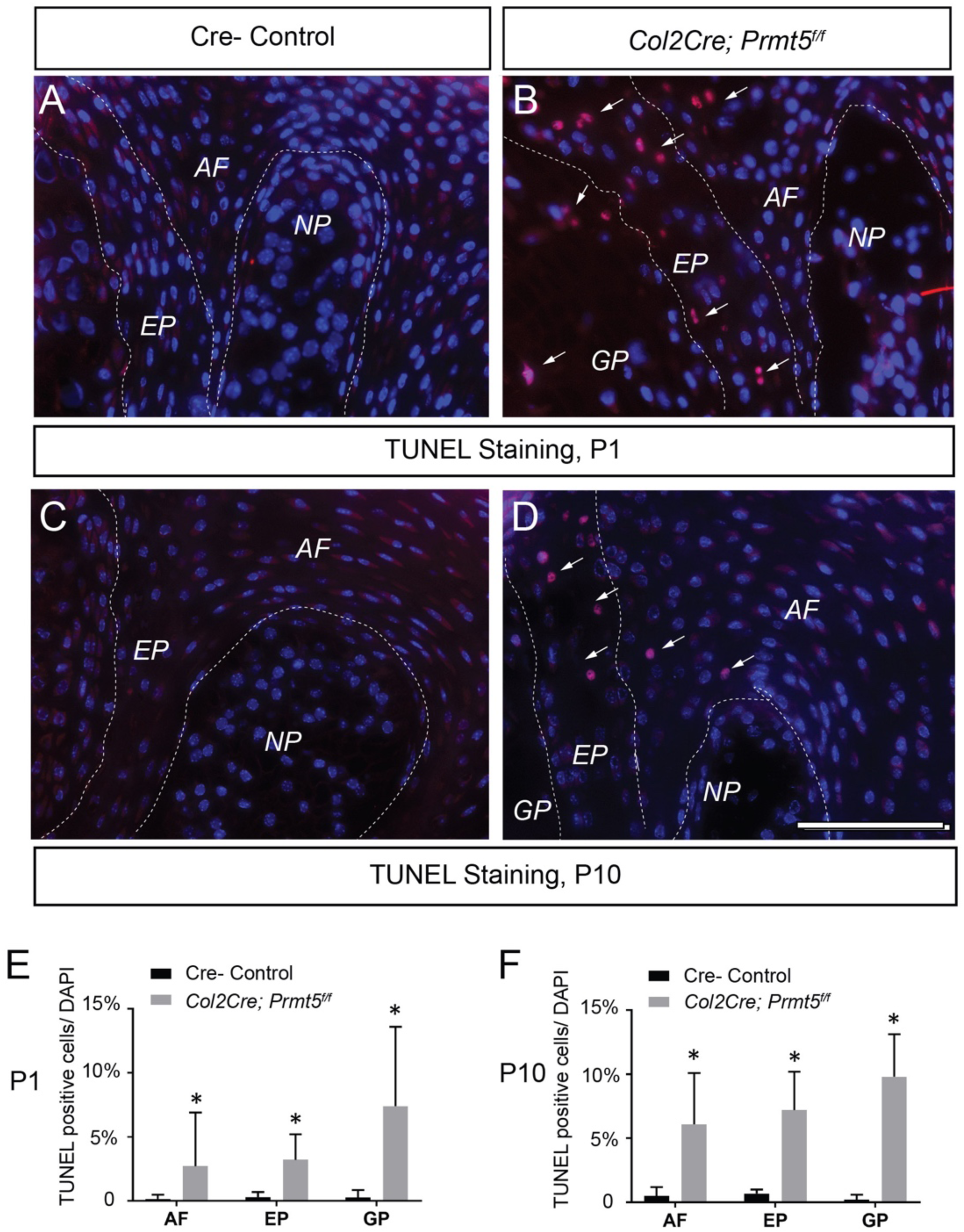
Loss of *Prmt5* in osteochondral progenitors results in increased cell death in the intervertebral disc. **(A-D)** TUNEL staining on spine sections of Cre (-) control (A, C) or *Col2Cre;Prmt5^f/f^* mutant (B, D) mice at P1 (A, B) and P10 (C, D). TUNEL positive cells were indicated with white arrows. (E, F) Quantification of TUNEL positive cells to DAPI positive cells in the IVD at P1 (E) and P10 (F). “*” indicates p<0.05, two-tailed student t-test. (*n*=3 for each group.) Scale bars: 100μm in (A-D). *AF: annulus fibrosus, EP: endplate, GP: growth plate, NP: nucleus pulposus*

**Supplementary Fig. 8.**
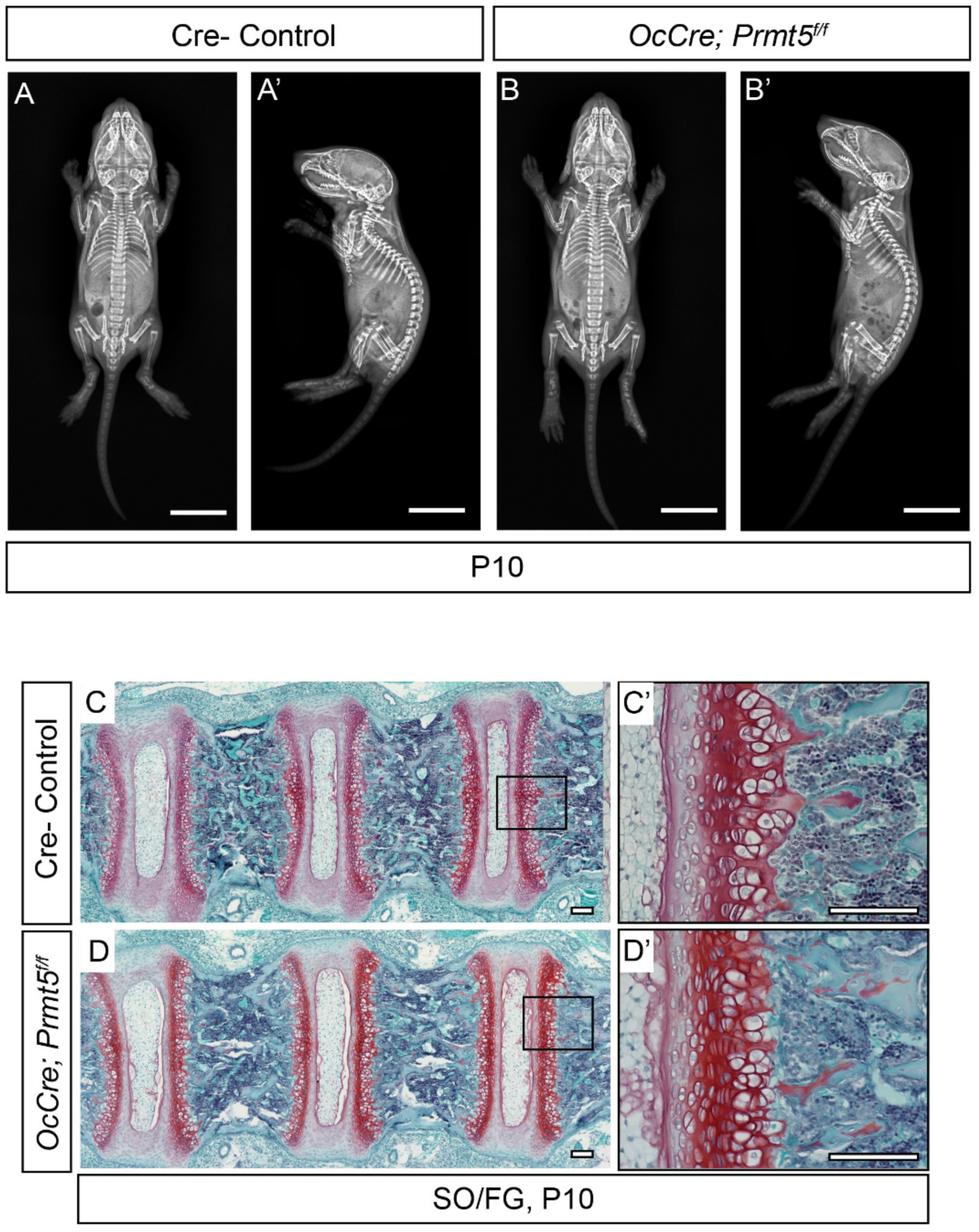
No scoliosis or spine deformity was observed in the *OcCre; Prmt5^f/f^* mice. **(A, B)** X-ray imaging analysis showed no in neither Cre scoliosis - control (A, A’) nor *OcCre; Prmt5^f/f^* mutant (B, B’) mice at P10. (C, D) Safranin O/Fast Green (SO/FG) staining on spine sections of Cre- control (C, C’) or *OcCre; Prmt5^f/f^* mutant mouse at P10. Images with higher magnification are indicated with black boxes and shown in C’ and D’. (*n*=4 for each group.) Scale bars: 10mm in (A-B’), 100μm in (C-D’).

**Supplemental Figure 9.**
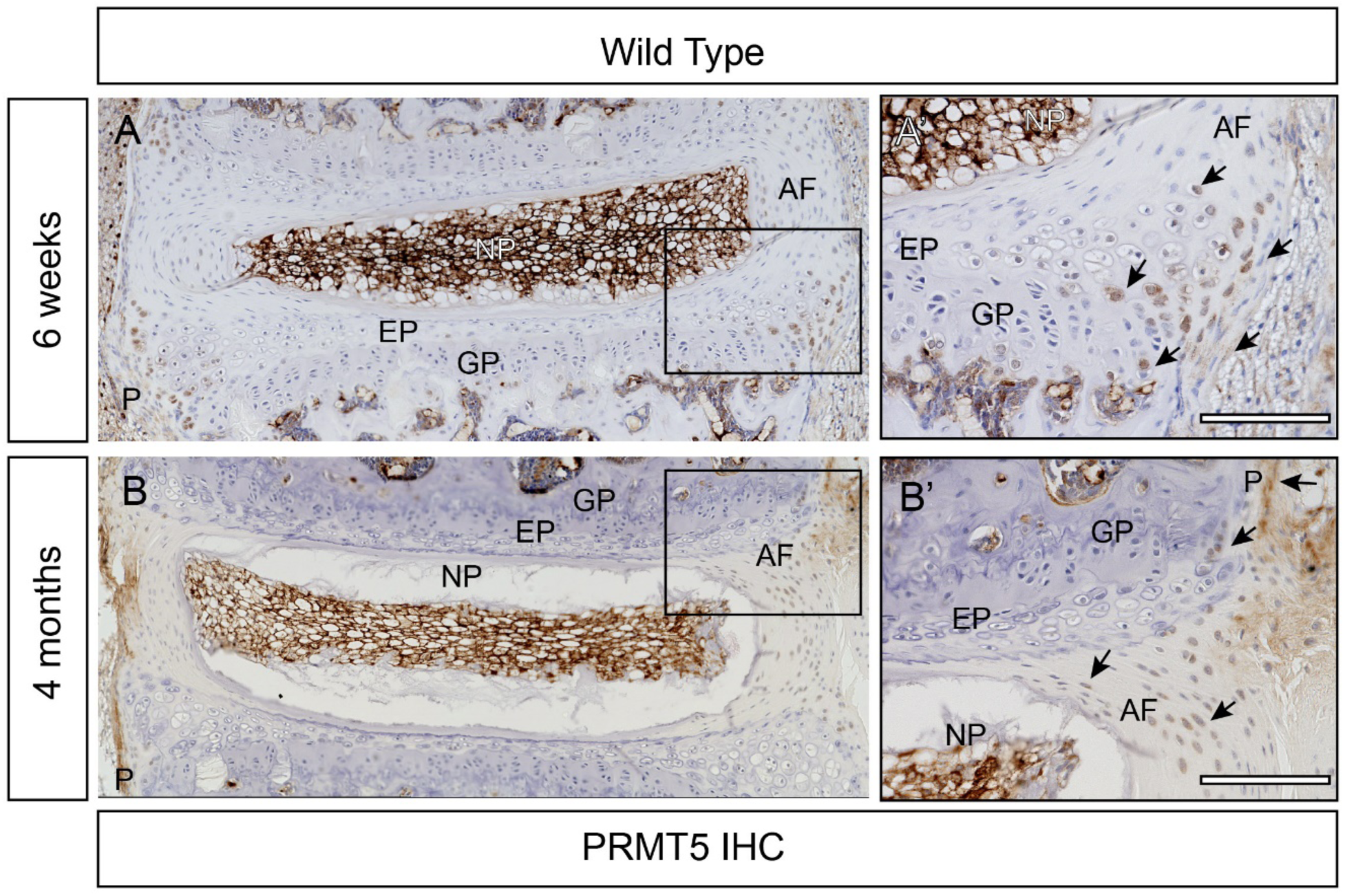
Low-level expression of PRMT5 was detected in postnatal IVD. **(A, B)** Immunohistochemistry (IHC) analysis of PRMT5 on spine sections of wild type mice at 6-weeks of age or 4-months of age, respectively. Images with higher magnification were indicated with black boxes and shown in A’, B’. Black arrows indicate PRMT5 positive cells. Scale bars: 100μm. *AF: annulus fibrosus, EP: endplate, GP: growth plate, NP: nucleus purposes, P: Perichondrium.* (*n*=3 for each group.)

**Supplemental Figure 10.**
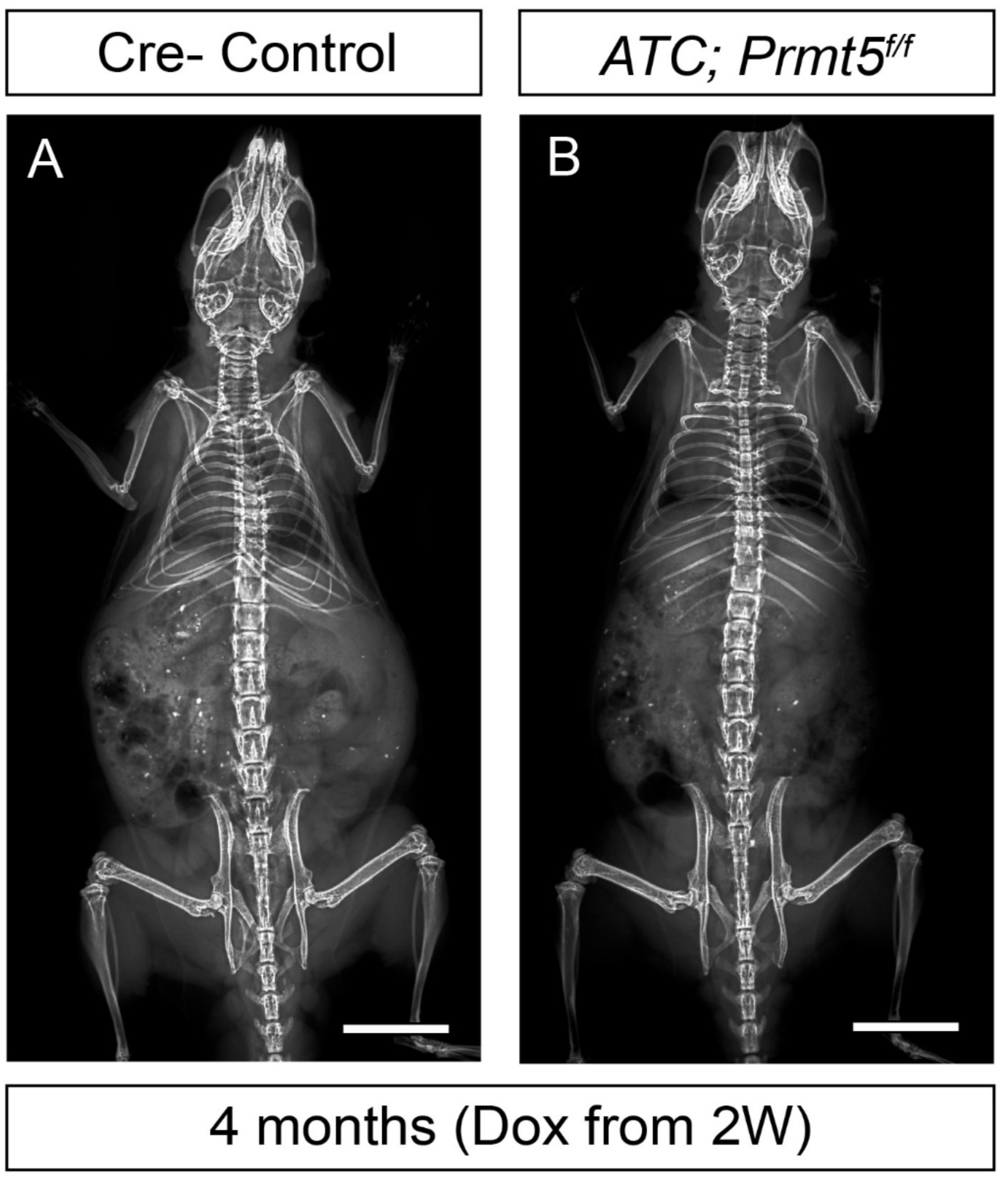
No scoliosis was observed in the *ATC; Prmt5^f/f^* mice when induced at 2-weeks of age. **(A, B)** X-ray imaging analysis showed no scoliosis in neither Cre- control (A) nor *ATC; Prmt5^f/f^* mutant (B) mice at 4-months of age (Dox induced from 2 weeks). (*n*=3 for each group.) Scale bars: 1mm.

